# Size and locomotor ecology have differing effects on the external and internal morphologies of squirrel (Rodentia: Sciuridae) limb bones

**DOI:** 10.1101/2023.02.08.527723

**Authors:** Johannah Rickman, Abigail E Burtner, Tate J Linden, Sharlene E Santana, Chris J Law

**Affiliations:** University of Washington

## Abstract

Mammals exhibit a diverse range of limb morphologies that are associated with different locomotor ecologies and structural mechanics. Much remains to be investigated, however, about the combined effects of locomotor modes and scaling on the external shape and structural properties of limb bones. Here, we used squirrels (Sciuridae) as a model clade to examine the effects of locomotor mode and scaling on the external shape and structure of the two major limb bones, the humerus and femur. We quantified humeral and femoral morphologies using 3D geometric morphometrics and bone structure analyses on a sample of 76 squirrel species across their four major ecotypes. We then used phylogenetic generalized linear models to test how locomotor ecology, size, and their interaction influenced morphological traits. We found that size and locomotor mode exhibit different relationships with the external shape and structure of the limb bones, and that these relationships differ between the humerus and femur. External shapes of the humerus and, to a lesser extent, the femur are best explained by locomotor ecology rather than by size, whereas structures of both bones are best explained by interactions between locomotor ecology and scaling. Interestingly, the statistical relationships between limb morphologies and ecotype were lost when accounting for phylogenetic relationships among species under Brownian motion. That assuming Brownian motion confounded these relationships is not surprising considering squirrel ecotypes are phylogenetically clustered; our results suggest that humeral and femoral variation partitioned early between clades and their ecomorphologies were maintained to the present. Overall, our results show how mechanical constraints, locomotor ecology, and evolutionary history may enact different pressures on the shape and structure of limb bones in mammals.

## Introduction

Mammals exhibit a variety of locomotor modes to transverse across a wide range of habitats (Hildebrand 1985). Adaptations to these various locomotor modes are repeatedly observed in mammalian limb bones (Polly 2007). For example, scansorial mammals tend to exhibit more elongate, gracile limb bones (Alexander et al. 1979; Burr et al. 1989; Kimura 1991; Polly 2007), whereas fossorial mammals exhibit relatively shorter and more robust limb bones in response to stress enacted on the bone when digging (Peterka 1936; Gasc et al. 1985; Straehl et al. 2013; Montoya-Sanhueza and Chinsamy 2017). Furthermore, different locomotor modes used in varying environments necessitate long bones being able to resist possible deformation to locomotion-specific forces. For example, species that fly, glide, or leap exhibit relatively long limbs with circular-shaped cross sections in the diaphyses to resist high torsional and multidimensional bending forces (Burr et al. 1989; Sharwtz et al. 1992; Patel et al. 2013; Hunt et al. 2021). In contrast, highly fossorial mammals exhibit large amounts of compact cancellous bone in the forelimb with elliptical-shaped cross sections to withstand uniaxial bending loads associated with digging (Amson et al. 2019; Amson et al. 2022).

Limb bone morphology is also influenced by body size (Alexander et al. 1979; Biewener 1983; Christiansen 1999). As body size increases, more mechanical support is needed to compensate for an increase in the mechanical forces exerted during locomotion (McMahon 1973; Biewener 1990; Christiansen 1999), which may result in adaptations in the shape and structural properties of the limb bones. Increasing body size is associated with increasing bone robustness in multiple lineages of tetrapods (Alexander et al. 1979; Demes and Jungers 1993; Christiansen 1999; Doube et al. 2009; Ryan and Shaw 2013; Mielke et al. 2018). In quadrupedal terrestrial tetrapods, body size rather than locomotor mode has a stronger influence on conserved limb bone structural traits such as the minor diaphyseal circumference of weight-bearing bones and the ratio between the stylopodial circumference and body mass (Campione and Evans 2012). Body size may also impose limits on bone shape; some cursorial carnivorans without the ability to supinate their forelimbs appear to have smaller maximum body sizes, while carnivorans with more generalized bone shapes exist at much larger body sizes (McMahon 1973; Polly 2007; Fabre et al. 2015).

Unsurprisingly, a plethora of work has examined the influence of locomotor behavior or body size on the overall shape of limb bones (e.g. Fabre et al. 2013; Martín-Serra et al. 2014a, 2014b; Hedrick et al. 2020; Etienne et al. 2021), and bone structural characteristics – such as bone compactness, diaphysis elongation, or cross-sectional shape – that are most directly impacted by different mechanical loads (e.g. Currey and Alexander 1985; Schaffler et al. 1985; Burr et al. 1989; Patel et al. 2013; Kilbourne and Hutchinson 2019; Scheidt et al. 2019; Wölfer et al. 2019; Amson et al. 2022). In this study, we further our understanding of limb bone morphological adaptations by testing how the interaction of locomotor ecology and scaling influences the shape and bone structure of the humerus and femur, using squirrels (Sciuridae) as a model clade. The squirrel family consists of approximately 280 species that inhabit a variety of microhabitats, from far above ground in tree canopies to deep underground in burrow systems. Squirrel species can be categorized into four ecotypes with distinct locomotor ecologies and behaviors: ground squirrels that dig, tree squirrels that climb, gliding squirrels that glide between trees, and more versatile chipmunks that both dig and climb. Previous work has examined relationships between limb lengths and ecotypes, finding, for example, that gliding squirrels exhibit relatively longer forelimbs than all other ecotypes (Peterka 1936; Bryant 1945; Thorington and Heaney 1981; Grossnickle et al. 2020; Linden et al. 2023), whereas ground squirrels exhibit relatively shorter forelimbs with increasing body elongation (Linden et al. 2023). Furthermore, across Sciuromorpha (squirrel-like rodents), cross-sectional characteristics (i.e., cross-sectional area and second moment of area) of the femur scale with positive allometry with respect to body mass (Scheidt et al. 2019). Nevertheless, whether the scaling patterns of external shape and bone structure of the humerus and femur differ among ecotypes remains to be tested within this clade. We predicted that humeral and femoral morphology is best explained by an interaction of ecotype and size, where both limb bones would be more robust, compact, and exhibit a more oval-shaped cross-section with increasing bone size in ground-dwelling (i.e., ground, tree, chipmunks) squirrels to resist the relatively greater mechanical loads that occur at larger body sizes (McMahon 1973; Biewener 1990; Christiansen 1999). Conversely, gliding squirrels would exhibit opposite trends. Within ecotypes, we predicted that ground squirrels would have more robust, compact limb bones with oval-shaped cross-sections to reinforce the bone in the cranial-caudal plane during scratch-digging (Hildebrand 1985, 1995; Lieberman et al. 2004; Lagaria and Youlatos 2006). We predicted that chipmunks and tree squirrels would exhibit more generalized limb bone shapes in comparison to other ecotypes, as they employ running, climbing, and digging behaviors. Finally, we predicted that gliding squirrels would exhibit more elongate, gracile limb bones with circular-shaped cross-sections to increase patagia surface area for gliding.

## Methods

### Morphological data

We acquired three-dimensional scans of humeri and femora from 76 squirrel species (one specimen per species) through computed tomography (CT) scanning with a Skyscan1172 μCT, Skyscan1173 μCT, GE Phoenix Nanotom M, and NSI X5000 scanning systems. Scans were performed with a resolution of 26.05μm. Scan data was reconstructed using NRecon and then exported and segmented in 3D Slicer (Kikinis et al. 2014). All specimens were sourced from the collections of 11 museums (Table S1). We used female, male, and sex-unknown individuals for our measurements to achieve the largest number of species. We determined that each specimen was fully mature by verifying that the cranial exoccipital-basioccipital and basisphenoid-basioccipital sutures were fused and that all vertebrae and limb bones were ossified.

### Ecotype classification

Following Linden et al. (2023), we categorized species into four ecotypes—chipmunk (n = 14), gliding (n = 8), ground (n = 25), or tree (n = 29)—based on locomotion, evolutionary grouping, and reproductive behavior (Fig. 1). Species that display fossorial locomotion and reproduce in underground burrows were categorized as ground squirrels, species that display both arboreal and scansorial locomotion and reproduce in nests in trees as were categorized tree squirrels, and species that have patagia were categorized as gliding squirrels. Our fourth ecotype group was chipmunks (genus *Tamias*), which display the broadest range of locomotor and nesting behaviors; species are considered terrestrial, semi-fossorial, or semi-arboreal depending on the source consulted, but none are considered fully fossorial or arboreal.

**Fig. 1.**
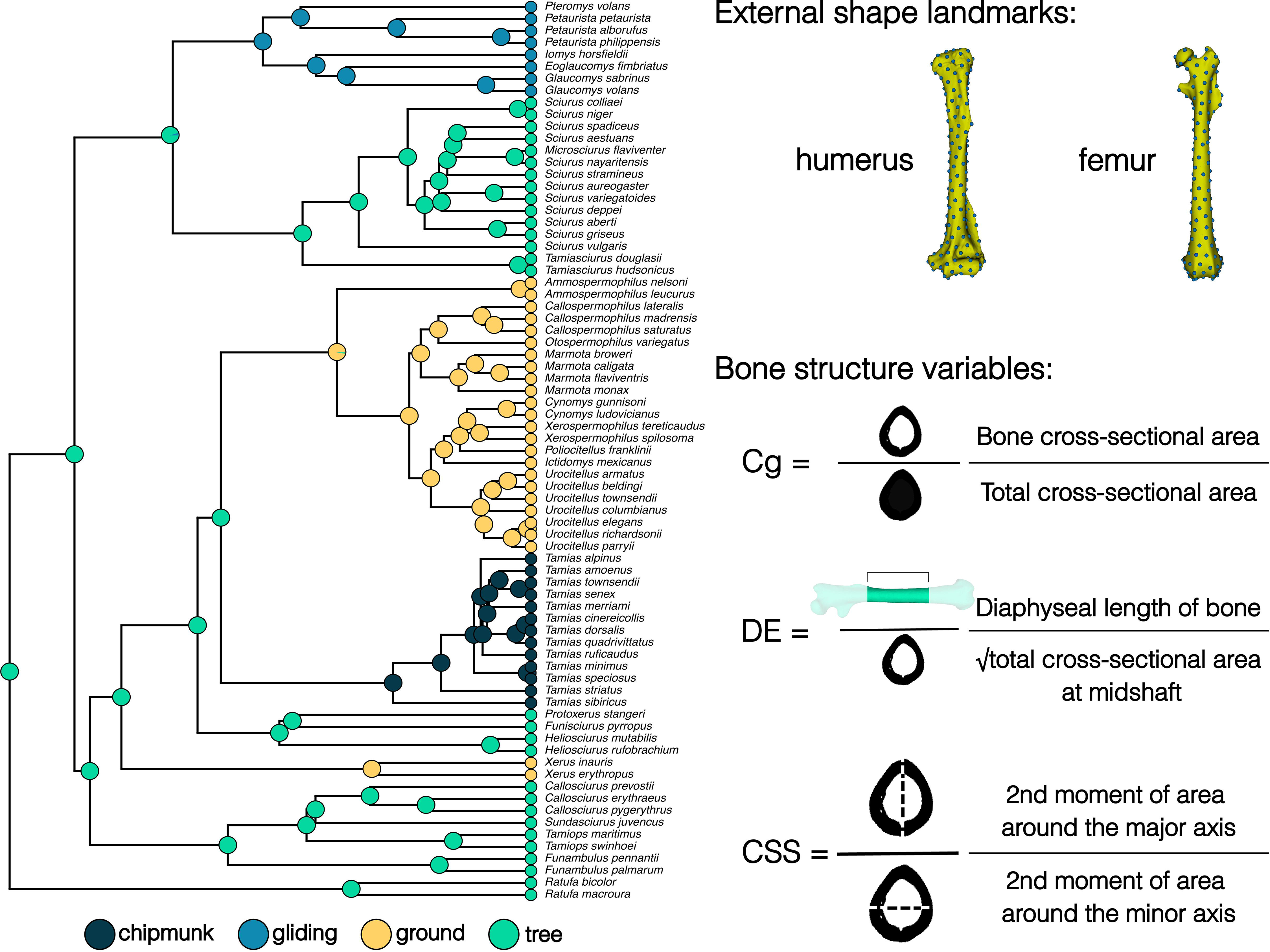
Overview of the species and morphometrics used. The pruned phylogeny is overlaid with an ancestral state reconstruction of squirrel ecotype. 194 pseudolandmarks on the humeri and 152 pseudolandmarks on the femora were generated and applied to our sample using the PseudoLMGenerator (Rolfe et al. 2021) and ALPACA modules (Porto et al. 2021) in SlicerMorph. Bone structure variables were calculated using the SegmentGeometry module (Huie et al. 2022) in Slicer. Cg = global compactness; DE = diaphysis elongation; CSS = cross-sectional shape.

### External shape analyses

All phylogenetic analyses were conducted using a pruned version of the Upham et al. (2019) phylogeny. All statistical analyses were performed in R (2022). We quantified humeral and femoral size and shape using three-dimensional geometric morphometrics (Rohlf and Slice 1990; Zelditch et al. 2012). We first transformed all specimens to correspond to the left side (an arbitrary choice) of the body using the Transforms module in 3D Slicer (Fedorov et al. 2012). We generated 194 pseudolandmarks on a humerus template and 152 pseudolandmarks on a femur template using the PseudoLMGenerator module of the SlicerMorph extension (Rolfe et al. 2021) in 3D Slicer (Fedorov et al. 2012). The most average squirrel in our dataset (determined visually on the phylomorphospace as the Western gray squirrel, *Sciurus griseus*) was used as our template. Following Diamond et al. (in press), we transferred the pseudolandmarks from the template model to each individual humerus and femur 3D model using the ALPACA module (Porto et al. 2021) in SlicerMorph (Fig. 1). We conducted a General Procrustes Analysis (GPA) using the gpagen function in the R package geomorph (Baken et al. 2021; Adams et al. 2022). We then verified that there were no significant shape differences between limb bones originating from the left or right side with Procrustes ANOVAs (humerus: F = 1.10, Z = 0.489, p = 0.319; femur: F = 1.38, Z = 1.03, p = 0.156).

We used humeral and femoral centroid size as our metrics of humeral and femoral size, respectively. We tested whether humeral and femoral size differed among ecotypes using phylogenetic analyses of variance (ANOVAs), which jointly estimate Pagel’s λ to account for phylogenetic covariance present in the model residuals, using the R package phylolm v2.6.2 (Tung Ho and Ané 2014). We assessed differences in limb sizes among ecotypes by first generating 1000 bootstrap replications of the model coefficients of limb size. We then computed the observed difference between the mean limb sizes of each ecotype pair and created its 95% confidence interval using the bootstrap replications. Confidence intervals that encompassed zero indicated that ecotype pairs were not significantly different from each other.

We visualized the phylomorphospace of humeral and femoral shape by performing principal component analyses (PCA) using the gm.prcomp function in geomorph (Baken et al. 2021; Adams et al. 2022). We also visualized differences between PC extremes by creating average shape models using the Morpho package in R (Schlager 2017) and used a custom Python script (Diamond et al. in press) to extract the mesh distances from the areas with pseudolandmark points.

We then tested how size, ecotype, and their interaction influenced humeral and femoral shape, respectively, using a series of phylogenetic penalized-likelihood multivariate generalized least squares (mvPGLS) models. These mvPGLS models use a penalized-likelihood to fit linear models to high-dimensional data sets in which the number of variables is much larger than the number of observations (Clavel et al. 2019). We fit our first model to test how each bone shape scaled with bone size (i.e., mvPGLS_size_ model: bone shape ∼ ln bone size). With the second model, we tested the relationship between bone shape and ecotype (i.e., mvPGLS_ecotype_ model: bone shape ∼ ecotype). With the third model, we tested if size and ecotype influenced bone shape (i.e., mvPGLS_size+ecotype_ model: bone shape ∼ ln bone size + ecotype). With the fourth model, we tested how the interaction of size and ecotype influenced bone shape (i.e., mvPGLS_size*ecotype_ model: bone shape ∼ ln bone size*ecotype). The last model we fit was a null model (mvPGLS_null_ model) in which size and ecotype have no influence on bone shape (i.e., bone shape ∼ 1). We assessed the importance of size, ecotype, their combined effect, and their interaction on limb shape variation by calculating the relative support for each of the five models through computation of Extended Information Criterion (EIC) weights (EICw) with 1000 bootstrap replications. We also assessed the scaling pattern of bone shape within each ecotype (i.e., bone shape_ecotype_ ∼ ln bone size_ecotype_). All mvPGLS models jointly estimated Pagel’s λ with the model parameters using the mvgls function (Clavel et al. 2019; Clavel and Morlon 2020) in the R package mvMORPH v.1.1.6 (Clavel et al. 2015). We performed the multivariate test using Pillai’s trace with 1000 permutations using the manova.gls function. We found that ecotype was an important predictor of bone shape (see Results); however, we were unable to perform post hoc pairwise testing among ecotypes because the mvMORPH function pairwise.glh is only suited for binary data thus far. Therefore, we performed post hoc pairwise permutation tests using the function pairwise in RRPP v1.3.1 (Collyer et al. 2015; Adams and Collyer 2018). These pairwise tests required us to repeat our analyses among ecotypes using Procrustes ANCOVAs in the R package geomorph v4.0.4 (Baken et al. 2021; Adams et al. 2022). Because our initial mvPGLS results indicated there was phylogenetic signal in the residuals of the model (Pagel’s λ = 0.18), we performed both (a) Procrustes ANCOVAs that assume no phylogenetic structure and (b) phylogenetic Procrustes ANCOVAs that assume Brownian motion to assess how the strength of phylogenetic signal in the residuals influenced our results. Procrustes ANCOVAs were performed with 1000 random residual permutation procedure (RRPP) using the functions procD.lm and procD.pgls in geomorph v4.0.4. Lastly, we estimated how well ecotype can predict limb shape using canonical variate analyses (CVA) with a jackknife cross-validation procedure in the R package Morpho (Schlager 2017).

### Bone structure analyses

We quantified three bone structural traits–global compactness (Cg), diaphysis elongation (DE), and cross-sectional shape (CSS)–that serve as proxies for biomechanical function (e.g., Patel et al. 2013; Hedrick et al. 2019; Kilbourne and Hutchinson 2019; Amson et al. 2021). Cg can reflect resistance towards axial compression where more compact limb bones (i.e., higher Cg) tend to resist more compressive loads (Berman et al. 2015). We used Cg instead of cross-sectional area because we aimed to quantify how much material is invested in the diaphysis of the limb bones while accounting for cross-section size. DE can be informative of the mechanical advantage of the limb movement, where relatively longer limb bones (i.e., higher DE values) reduce the out-lever whereas relatively shorter limb bones (i.e., lower DE values) increase the out-lever (Smith and Savage 1956). CSS can reflect torsion and uni- or multi-directional bending of the humeral and femoral diaphyses, where bones with more elliptical cross-sections (i.e., higher CSS values) are hypothesized to be adapted for bending loads in one direction, whereas bones with more circular cross-sections (i.e., lower CSS values) are hypothesized to be adapted for the multidirectional bending loads that are associated with using diverse locomotor behaviors or resisting torsion during aerial locomotion (Sharwtz et al. 1992; Patel et al. 2013; Amson et al. 2021). We quantified these three traits by creating slice-by-slice proximodistal profiles of each scanned specimen in the Slicer module SegmentGeometry (Huie et al. 2022). Each profile consisted of length (in millimeters), cross-sectional area (in square millimeters), second moment of area around the major principal axis, second moment of area around the minor principal axis, and compactness (ratio between sample cross-sectional area and total cross-sectional area). Outputs from SegmentGeometry were used to calculate three bone structure traits: Cg, defined as the mean compactness of all included slices; DE, defined as the ratio between the functional length of the bone and the square-root of the cross-sectional area at the midshaft slice; and CSS, defined as the ratio of the second moment of area around the major principal axis to the second moment of area around the minor principal axis at the midshaft slice (Fig. 1). Each bone structure trait was computed across the middle 40% of the bones’ functional length, which captures the greatest distance between the proximal and distal articular surfaces of the bones and ensures only the diaphysis is measured.

We tested how size, ecotype, and their interaction influenced each bone structure trait (i.e., Cg, DE, CSS) using a series of phylogenetic generalized least squares (PGLS) models. Similar as above, we assessed the importance of size, ecotype, their combined effect, and their interaction on bone structure traits by fitting five models (i.e., PGLS_size_, PGLS_ecotype_, PGLS_size+ecotype_, PGLS_size*ecotype_, PGLS_null_) and calculating the relative support for each model through computation of Akaike information criterion weights (AICw). All models with ΔAIC below 2 were considered to be supported by the data (Burnham and Anderson 2002). Regression coefficients for all models were estimated simultaneously with phylogenetic signal as Pagel’s λ in the residual error using the R package phylolm v2.6.2 (Tung Ho and Ané 2014). We also generated 1000 bootstrap replications of the scaling slopes and means of each variable in each PGLS model. In instances where the best fitting model incorporated ecotype, we assessed if bone structure variables differed among ecotypes by computing the observed difference between the mean values of each ecotype pair and creating its 95% confidence interval using the bootstrap replications. Confidence intervals that encompassed zero indicated that ecotype pairs were not significantly different from each other. To maintain consistency with the external shape analyses, we also assessed if bone structure variables differed among ecotypes using (a) ANCOVAs that assume no phylogenetic structure and (b) phylogenetic ANCOVAs that assume full Brownian motion. All ANCOVAs were performed with 1000 RRPP iterations using the function lm.rrpp in the R package RRPP (Collyer and Adams 2018). In instances where the best fitting model incorporated size, we determined if bone structure scaling was significantly different from isometry across all squirrels. To do this, we used the bootstrap replications to create 95% confidence intervals for the mean scaling slopes across all ecotypes as well as for each individual ecotype. Confidence intervals greater than zero were considered to represent positive allometry, while confidence intervals less than zero were considered to represent negative allometry. Confidence intervals that included zero were considered isometric due to the dimensionless quality of the bone structure variables. In instances where the best fitting model incorporated the interaction between ecotype and size, we assessed if scaling patterns of bone structure variables differed among ecotypes by computing the observed difference between the mean slopes of each ecotype pair and created its 95% confidence interval using the bootstrap replications. Confidence intervals that encompassed zero indicated that ecotype pairs were not significantly different from each other. We also assessed if bone structure scaling was significantly different from isometry within each ecotype by using the bootstrap replications to create 95% confidence intervals for the mean scaling slopes for each individual ecotype. Confidence intervals greater than zero were considered to represent positive allometry, while confidence intervals less than zero were considered to represent negative allometry. Confidence intervals that included zero were considered isometric due to the dimensionless quality of the bone structure variables.

### Comparisons between bone shape and structure, and between humeral and femoral traits

We tested if there were relationships between the external shape and each of the bone structure variables of the humerus and femur across squirrels and within each ecotype using two-block partial least squares (PLS) analyses in the R package geomorph version 4.0.4 (Baken et al. 2021; Adams et al. 2022). We also tested if there were relationships between humeral traits and femoral traits across all squirrels and within each ecotype. We used PLS analyses in geomorph to test relationships between humeral shape and femoral shape, and phylogenetic paired t-tests in the phytools function phyl.pairedttest to test if each bone structure variable differed between the humerus and femur.

## Results

### Humeral and femoral size

We found that ecotype explained 4% of humeral size variance (R^2^ = 0.04; Pagel’s λ = 0.95). Pairwise comparisons between mean humeral sizes revealed that chipmunks (mean humeral size [95% CI] = 4.6 [3.8:5.3] cm) exhibited the smallest humeri, followed by tree squirrels (5.2 [4.8:5.7] cm), gliding squirrels (5.4 [4.8:6.1] cm), and ground squirrels (5.0 [4.4:5.6] cm) (Fig. 2A; Table S2).

**Fig. 2.**
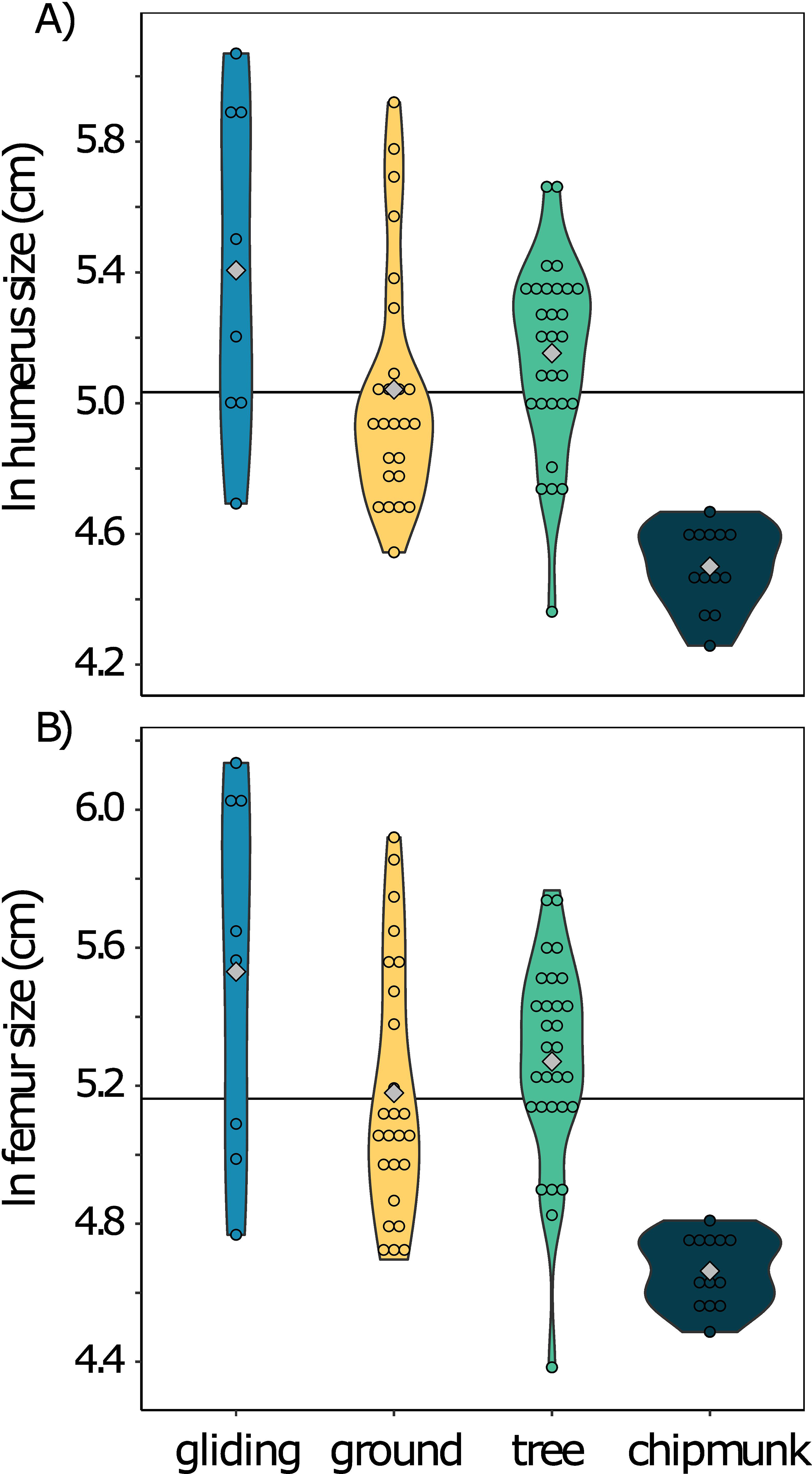
Violin plots of ln humeral size and ln femoral size by ecotype. The gray diamonds indicate the mean of each ecotype, and the horizontal black line indicates the mean size across all ecotypes.

**Fig. 3.**
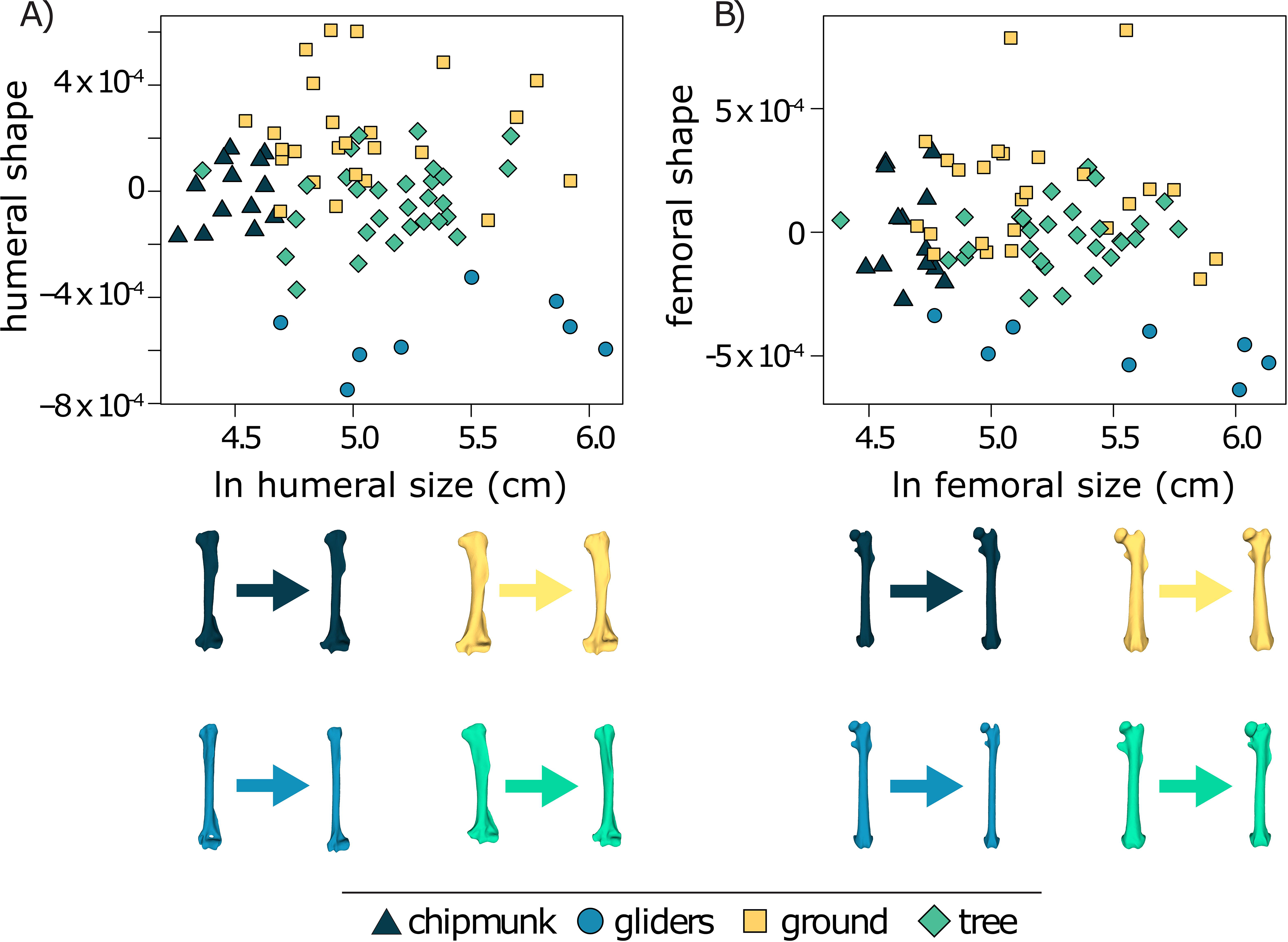
Allometric trends between external shape and size of the A) humerus and B) femur. Limb bone images show the difference in shape from the smallest to biggest species in each ecotype (chipmunk: *Tamias alpinus* to *Tamias townsendii*; gliding: *Glaucomys volans* to *Petaurista petaurista*; ground: *Callospermophilus madrensis* to *Marmota caligata*; tree: *Tamiops maritimus* to *Ratufa macroura*).

We found that ecotype explained 3% of femoral size variance (R^2^ = 0.03; Pagel’s λ = 0.96). Pairwise comparisons between mean femoral sizes revealed that chipmunks (mean femoral size [95% CI] = 4.8 cm [4.0:5.6 cm]) exhibited the smallest femora, followed by ground squirrels (5.2 cm [4.6:5.9 cm]), tree squirrels (5.4 [4.9:5.8 cm]), and gliding squirrels (5.6 cm [4.8:6.3 cm]) (Fig. 2B; Table S2).

### Humeral external shape

PC1 accounted for 36.8% of the humeral shape variation and largely separates the morphospaces among ground squirrels, chipmunks and tree squirrels, and gliding squirrels (Fig. 4). PC1 primarily describes the lengthening of the humerus. Positive PC1 scores are associated with more robust humeri with more pronounced supinator crests, deltoid tuberosities, and medial epicondyles, whereas negative PC1 scores are associated with more elongate humeri with less developed supinator crests, deltoid tuberosities, and medial epicondyles. PC2 accounted for 12.5% of the humeral shape variation and is associated with shape variation at the ends of the humerus (Fig. 4). Positive PC2 scores are associated with shape variation at the medial epicondyle and negative PC2 scores are associated with changes at the condyle. Much of this variation across PC2 explains within-ecotype differences.

**Fig. 4.**
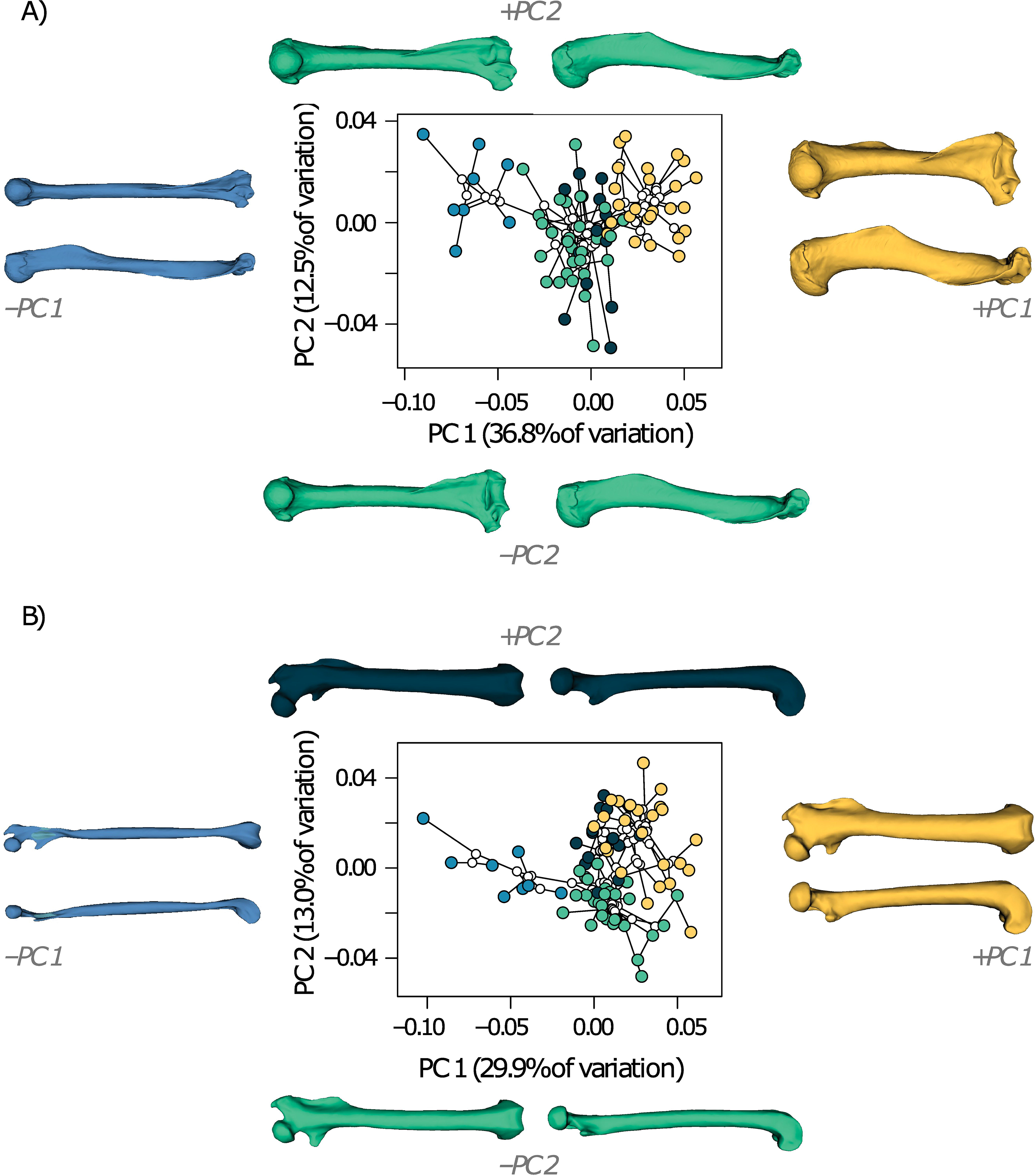
Phylomorphospaces of A) humeral shape and B) femoral shape with PC1-2 extremes for visualization. The humeral and femoral models reflect the computationally determined PC1 and 2 extremes, and their colors reflect the ecotype associated with extreme ends of the first two PCs. Blue circles represent gliding squirrels, yellow circles represent ground squirrels, navy circles represent chipmunks, green circles represent tree squirrels, and open circles represent tree nodes within the phylomorphospace.

The mvPGLS_ecotype_ model was the best supported model for humeral shape (EICw = 1.00; Table 1) and showed a statistically significant relationship between humeral shape and ecotype (Pagel’s λ = 0.19, Pillai’s trace = 2.83, P < 0.001). Because we were unable to perform post hoc pairwise tests among ecotypes using the PL-MANOVA model (see Methods), we performed pairwise permutation tests using both phylogenetic Procrustes ANCOVA that assumes Brownian motion and Procrustes ANCOVA that does not incorporate phylogenetic structure. Unlike the mvPGLS_ecotype_ model, the phylogenetic Procrustes ANCOVA model indicated that there was no significant relationship between humeral shape and ecotype (R^2^ = 0.04, Z = -0.137, P = 0.550). In contrast, the non-phylogenetic Procrustes ANCOVA model indicated that there was a significant relationship between humeral shape and ecotype (R^2^ = 0.37, Z = 6.551, P < 0.001), and post-hoc pairwise tests revealed that there were significant differences in humeral shape among all ecotypes (all P < 0.002) except between chipmunks and tree squirrels (P = 0.075). This is consistent with the CVA with jackknife cross-validation, which reclassified humeral shapes in their correct ecotype with 93.4% accuracy overall and accurately reclassified 100% of gliding and ground squirrels, 85.7% of chipmunks, and 89.7% of tree squirrels.

**Table 1.**
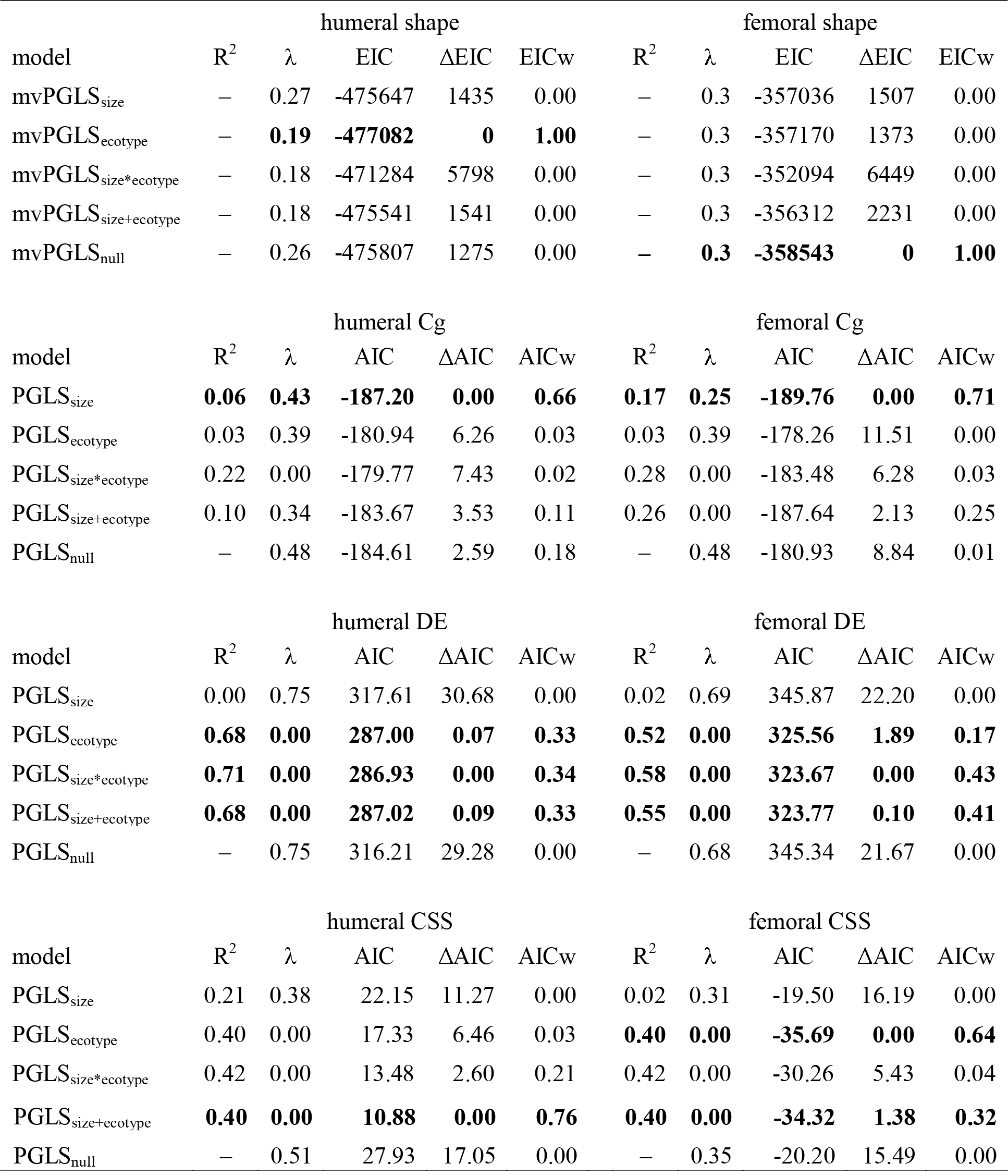
Comparisons of the best-fitting phylogenetic generalized least squares (PGLS) models in humeral and femoral shape and bone structure. Extended Information Criterion (EIC) and Akaike Information Criterion (AICc) were used to assess model fits on bone shape and bone structure, respectively. Rows in boldface type represent the best-fit model as indicated by the lowest ΔEIC or ΔAIC score. λ = Pagel’s λ; EICw = EIC weights; AICw = AIC weights.

Within ecotypes, gliding squirrels (Pagel’s λ = 0.00, Pillai’s trace = 1.00, P = 0.016) and ground squirrels (Pagel’s λ = 0.21, Pillai’s trace = 0.98, P = 0.009) exhibited significant humeral shape allometry whereas chipmunks (Pagel’s λ = 0.00, Pillai’s trace = 0.98, P = 0.382) and tree squirrels (Pagel’s λ = 0.26, Pillai’s trace = 0.94, P = 0.742) did not.

### Femoral external shape

PC1 accounted for 29.3% of the femoral shape variation and separated the morphospace of gliding squirrels from a morphospace largely containing ground squirrels, chipmunks, and tree squirrels (Fig. 4). Positive PC1 scores are associated with more robust femora with more pronounced femoral heads, patellar grooves, greater trochanters, third trochanters, and lesser trochanters whereas negative PC1 scores are associated with more elongate, gracile femora (Fig. 4). PC2 accounted for 12.8% of the femoral variation and loosely separated tree squirrels from ground squirrels and chipmunks. Positive PC2 scores are associated with shape variation around the greater trochanter and one side of the lesser trochanter, whereas negative PC2 scores are associated with shape variation around the femoral head, greater trochanter, and the opposite side of the lesser trochanter (i.e., one extreme reflects shape variation “bending” the trochanter out and the other in dorsal-ventrally) (Fig. 4).

The mvPGLS_null_ model was the best supported model for femoral shape (EICw = 1.00; Table 1), suggesting that neither size nor ecotype are important predictors of femoral shape. Nevertheless, the CVA accurately reclassified femoral shapes in their correct ecotype with 90.8% accuracy overall and accurately reclassified 100% of gliding squirrels, 96.6% of tree squirrels, 92.9% of chipmunks, and 80.0% of ground squirrels. Consistent with the CVA, results from the Procrustes ANCOVA model that does not incorporate phylogenetic structure indicated that there was a significant relationship between femoral shape and ecotype (R^2^ = 0.31, Z = 6.88, P < 0.001), and post-hoc pairwise tests revealed that there were significant differences in femoral shape among all ecotypes (all P < 0.004). However, incorporating phylogenetic structure resulted in no significant relationship between femoral shape and ecotype (mvPGLS_ecotype_ model with Pagel’s λ = 0.32: Pillai’s trace = 2.57, P = 0.194; phylogenetic Procrustes ANCOVA under Brownian motion: R^2^ = 0.04, Z = -0.446, P = 0.676).

Within ecotypes, gliding squirrels (Pagel’s λ = 0.00, Pillai’s trace = 0.99, P = 0.024) and tree squirrels (Pagel’s λ = 0.26, Pillai’s trace = 0.97, P = 0.008) exhibited significant femoral shape allometry, whereas chipmunks (Pagel’s λ = 0.00, Pillai’s trace = 0.96, P = 0.823) and ground squirrels (Pagel’s λ = 0.50, Pillai’s trace = 0.96, P = 0.071) did not.

### Humeral bone structure

Size is a more important predictor of humeral Cg than ecotype, as the PGLS_size_ model received the greatest support (R^2^ = 0.06, Pagel’s λ = 0.43, AICw = 0.66; Table 1). We found that, across all squirrels, humeral Cg was positively allometric relative to humeral size (slope [95% CI] = 0.05 [0.00:0.10]), indicating squirrels evolved more compact humeri as humeral size increased. Within ecotypes, only tree squirrel (0.12 [0.03:0.19]) humeral Cg exhibited positive allometry, whereas chipmunk (-0.15 [-0.14:0.44]), ground squirrel (0.03 [-0.04:0.10]) and gliding squirrel (0.04 [-0.05:0.13]) humeral exhibited isometry.

Size, ecotype, and their interaction are important predictors of humeral DE, with the PGLS_size*ecotype_ model (R^2^ = 0.71, λ = 0.00, AICw = 0.34), the PGLS_ecotype_ (R^2^ = 0.68, λ = 0.00, AICw = 0.33), and the PGLS_size+ecotype_ (R^2^ = 0.68, λ = 0.00, AICw = 0.33) models receiving equal support (Table 1). Scaling slopes differed among ecotypes (Table S2). Tree squirrel humeral DE exhibited negative allometry (-2.07 [-3.88:-0.29]), indicating that tree squirrels evolved less slender humeri as humeral size increased. Chipmunk (0.05 [-6.42:6.70), gliding squirrel (1.42 [- 0.61:3.44), and ground squirrel (-1.10 [-2.55:0.48]) humeral DE exhibited isometry. Comparisons of the differences between mean humeral DE values among ecotypes indicate that gliding squirrels tend to exhibit the slenderest humeral diaphyses (mean humeral DE [95% CI] = 22.39 [21.27:23.51]), followed by chipmunks (17.32 [16.44:18.14]), tree squirrels (16.92 [16.38:17.47]), and ground squirrels (14.81 [14.19:15.41]), Fig. 6; Table S2). Differences in humeral DE among ecotypes are corroborated by with an ANOVA (R^2^ = 0.68, P = 0.001; all pairwise comparisons P < 0.009 except for the chipmunk-tree comparison) but not the phylogenetic ANOVA (R^2^ = 0.04, P = 0.356). Humeral DE was isometric relative to humeral size across all ecotypes (slope [95% CI] = -0.60 [-2.11:0.89]).

Both size and ecotype are important predictors of humeral CSS, as the PGLS_size+ecotype_ model received the greatest support (R^2^ = 0.40, Pagel’s λ = 0.43, AICw = 0.76; Table 1). Across all squirrels, humeral CSS exhibited positive allometry (slope [95% CI] = 0.28 [0.08 - 0.47]), which indicates that squirrels evolved more oval-shaped humeral cross-sections as humeral size increased. Within ecotypes, ground squirrel humeral CSS exhibited positive allometry with humeral centroid size (0.42 [0.18 - 0.64]), whereas chipmunks (0.37 [-0.64:1.46]), gliding squirrels (0.02 [-0.32:0.38]), and tree squirrels (0.21 [-0.08:0.49]) exhibited isometry. Comparisons of the differences between mean humeral CSS values among ecotypes indicate that ground squirrels tend to exhibit the most oval-shaped humeral cross-sections (mean humeral CSS [95% CI] = 1.71 [1.61:1.81]) followed by tree squirrels (1.45 [1.36:1.54]). Chipmunks (1.26 [1.13:1.39]) and gliding squirrels (1.26 [1.09:1.43]) tend to exhibit similarly low CSS values (Fig. 6; Table S2). Differences in humeral CSS among ecotypes are corroborated by with an ANOVA (R^2^ = 0.33, P = 0.001); pairwise tests showed ground squirrels had more oval-shaped humeral cross-sections compared to all other ecotypes (all P < 0.004) whereas the remaining ecotypes exhibited no differences in humeral CSS (all P > 0.071). In contrast, the phylogenetic ANOVA indicated no significant relationship between humeral CSS and ecotype (R^2^ = 0.01, P = 0.839).

### Femoral bone structure

Size is a more important predictor of femoral Cg than ecotype, with PGLS_size_ being the best-supported model (R^2^ = 0.17, λ = 0.25, AICw = 0.71; Table 1). Across all squirrels, femoral Cg was positively allometric (slope [95% CI] = 0.08 [0.04:0.13]), indicating that squirrels evolved more compact femora as femoral size increased. Within ecotypes, ground squirrel (0.08 [0.01:0.14]) and tree squirrel (0.12 [0.05:0.20]) and femoral Cg exhibited positive allometry, whereas chipmunk (-0.05 [-0.37:0.28]) and gliding squirrel (0.06 [-0.03:0.14]) femoral Cg exhibited isometry.

Size, ecotype, and their interaction are important predictors of femoral DE; there was equal support for the PGLS_size*ecotype_ (R^2^ = 0.58, λ = 0.00, AICw = 0.43) and the PGLS_size+ecotype_ (R^2^ = 0.55, λ = 0.00, AICw = 0.41) models (Table 1). Across all squirrels, femoral DE was isometric relative to femoral size (slope [95% CI] = -1.08 [-2.97:0.81]). Within ecotypes, tree squirrel (-2.22 [-4.54:-0.02]) and ground squirrel (-2.41 [-4.38:-0.43]) femoral DE exhibited negative allometry, indicating that tree and ground squirrels tend to evolve less slender femora as femoral size increases. Chipmunk (-1.99 [-12.54:7.70]) and gliding squirrel (1.51 [-0.91:3.93]) femoral DE exhibited isometry. Comparisons of the differences between mean femoral DE values among ecotypes indicate that gliding squirrels tend to exhibit the slenderest femoral diaphyses (mean femoral DE [95% CI] = 25.00 [23.58:26.42]) whereas ground squirrels tend to exhibit the most robust femoral diaphyses (18.02 [17.26:18.76]). Chipmunks (21.05 [20.00:22.12]) and tree squirrels (20.40 [19.70:21.10]) tend to exhibit similar femoral DE values (Fig. 6; Table S2). Mean differences in femoral DE among ecotypes were corroborated by the ANOVA (R^2^ = 0.52, P = 0.001; all pairwise comparisons P < 0.003 except for the chipmunk-tree comparison) but not the phylogenetic ANOVA (R^2^ = 0.03, P = 0.593).

Ecotype is a more important predictor of femoral CSS than size, as PGLS_ecotype_ was the best supported model (R^2^ = 0.40, λ = 0.00, AICw = 0.64; Table 1). There were only slight differences of mean femoral CSS values between ground squirrels (mean femoral CSS [95% CI] = 1.91 [1.23:2.56]), chipmunks (1.80 [1.21:2.39]), tree squirrels (1.63 [0.96:2.29]), and gliding squirrels (1.55 [0.85:2.26]) (Fig. 6; Table S2). Mean differences in femoral CSS among ecotypes were corroborated by the ANOVA (R^2^ = 0.40, P = 0.001; all pairwise comparisons P < 0.006 except for the chipmunk-ground and gliding-tree comparisons) but not the phylogenetic ANOVA (R^2^ = 0.01, P = 0.834).

### Relationships between external shape and bone structure

We found that humeral shape exhibited significant relationships with DE (r-PLS = 0.88, Z = 4.547, P < 0.001) and CSS (r-PLS = 0.62, Z = 3.257, P < 0.001) but not Cg (r-PLS = 0.39, Z = 0.956, P = 0.186; Table S3). Within ecotypes, only tree squirrel humeral shape exhibited significant relationships with all three bone structure traits, and ground squirrel humeral shape exhibited a significant relationship with humeral DE and humeral CSS (Table S3).

We found that femoral shape exhibited significant relationships with Cg (r-PLS = 0.58, Z = 2.813, P < 0.001) and DE (r-PLS = 0.85, Z = 4.702, P < 0.001), and CSS (r-PLS = 0.63, Z = 3.133, P < 0.001). Within ecotypes, ground squirrel femoral shape exhibited a significant relationship with femoral Cg and femoral DE, and tree squirrel femoral shape exhibited a significant relationship with femoral DE (Table S3).

### Comparisons between humeral and femoral shape and bone structure

We found a significant relationship between humeral shape and femoral shape across all squirrels (r-PLS = 0.91, Z = 5.884, P < 0.001). Within ecotypes, gliding squirrels (r-PLS = 0.98, Z = 2.519, P = 0.004) and tree squirrels (r-PLS = 0.83, Z = 2.236, P = 0.012) exhibited a significant relationship between humeral shape and femoral shape, but neither chipmunks (r-PLS = 0.81, Z = 0.672, P = 0.281) nor ground squirrels (r-PLS = 0.82, Z = 1.523, P = 0.068) did.

We found that global compactness (Cg) differed between the humerus (mean humeral Cg = 0.63 and femur (mean femoral Cg = 0.57) across all squirrels (phylogenetic mean difference = 0.05, P < 0.001) as well as within each ecotype (all P < 0.043) except gliding squirrels (P = 0.595; Table S4). Diaphysis elongation (DE) also differed between the humerus (mean humeral DE = 16.87) and femur (mean femoral DE = 20.21) across all squirrels (phylogenetic mean difference = 3.34, P < 0.001) and within each ecotype (all P < 0.001; Table S4). Lastly, there was no significant difference in cross-sectional shape (CSS) between the humerus (mean humeral CSS = 1.48) and femur (mean femoral CSS = 1.48) across all squirrels (P = 0.530) and within each ecotype (all P > 0.065) except in chipmunks (P < 0.001; Table S4).

## Discussion

Size, locomotor ecology, and phylogenetic history are often identified as the leading factors that underlie limb bone variation in mammals (e.g., Fabre et al. 2013; Kilbourne and Hoffman 2013; Martín-Serra et al. 2014b, 2014a; Mielke et al. 2018; Scheidt et al. 2019; Wölfer et al. 2019; Etienne et al. 2021). Here, we found that these factors exhibit different relationships with the external shape and structure of squirrel humeri and femora: ecotype influenced variation of the external shape of these bones, whereas interactions between size and ecotype influenced variation of their structure.

Our results suggest that differences in locomotor behavior have a stronger effect on humeral shape evolution and, to a lesser extent, femoral shape evolution rather than size (contrary to our predictions; Table 1). Using our model selection approach, we found that humeral shape is best described by the mvPGLS_ecotype_ model (Table 1), which revealed a significant relationship between humeral shape and ecotype. Gliding squirrels tend to exhibit gracile, elongate humeri; ground squirrels tend to exhibit robust humeri; and chipmunks and tree squirrels tend to exhibit intermediate humeri (Fig. 4; Table 1). However, the statistical relationship between ecotype and humeral shape was lost when using a model that assumes Brownian motion. In contrast, we found that femoral shape is best described by the mvPGLS_null_ model (Table 1), suggesting that femoral shape is not well predicted by ecotype, size, or their interaction. Nevertheless, the CVA indicated that femoral shape can be accurately reclassified by ecotype, and the Procrustes ANOVA also supported a significant relationship between femoral shape and ecotype. However, like in the humeral shape analyses, the statistical relationship between femoral shape and ecotype is lost when incorporating phylogenetic signal as Pagel’s λ or full Brownian motion (i.e., λ = 1). That accounting for phylogenetic relationships confounded these relationships is not surprising considering squirrel ecotypes are phylogenetically clustered (Fig. 4); chipmunks and gliding squirrels are each monophyletic, ground squirrels consist of two clades, and tree squirrels are ancestral (Fig. 1). These results also indicate that there was no evolutionary transition from relatively elongate, gracile humeri and femora to relatively shorter, robust ones with increasing size across squirrels, a pattern that is found in other mammalian clades (e.g., Fabre et al. 2013; Martín-Serra et al. 2014a, 2014b; Etienne et al. 2021). Instead, the lack of shape allometry in the proximal limb bones is consistent with what is found in myomorph and geomyoid rodents (Hedrick et al. 2019). This provides additional evidence that only large mammals require major allometry-driven shape changes (Biewener 2005) or allometric changes occur in other aspects of the limb morphology such as bone structure (see next section). Overall, our results emphasize that locomotor behavior has a stronger influence on humeral shape compared to femoral shape, and that the diversity of squirrel humeral shapes and to a lesser extent, femoral shapes, were ecologically partitioned early between clades and these ecomorphologies were maintained to the present.

In contrast, bone structure variables of the humerus and femur exhibited various scaling trends across all squirrels. Global compactness (Cg) of both the humerus and femur was best described by the PGLS_size_ model (Table 1), which shows Cg increased with increasing limb size (Fig. 5). This pattern is consistent with findings that cortical bone thickness (Currey and Alexander 1985) and cross-sectional properties of the femur (Mielke et al. 2018; Scheidt et al. 2019) scale with increasing size. These positive allometric relationships may be due to the increase in mechanical stress at larger body sizes (McMahon 1973; Biewener 1990; Christiansen 1999). Positive allometry towards more compact bones may also free the constraints of allometry-driven changes in limb bone shape; the mechanical demands against gravity may be met with positive allometry of Cg, enabling other ecological factors to have greater influence on limb bone shapes. Cross-sectional shape (CSS) of the humerus and femur were best described by models that included both size and ecotype (Table 1). These models revealed that scaling patterns differed between humeral CSS and femoral CSS; humeral cross-sections became more oval-shaped as size increased through positive allometry, whereas femoral CSS exhibited isometry. Diaphysis elongation (DE) of both the humerus and femur were best described by models that included both size and ecotype and their interaction (Table 1). These models revealed that scaling patterns of humeral and femoral DE did not statistically differ from isometry across all squirrels. Like the bone shape analyses, the statistical relationships between bone structure variables and ecotype were lost when using phylogenetic ANOVAs that assume Brownian motion but not with the non-phylogenetic ANOVAs. These results are consistent with the bone shape analyses, providing additional support that humeral and femoral morphologies were ecologically partitioned early between clades and their ecomorphologies were maintained to the present.

**Fig. 5.**
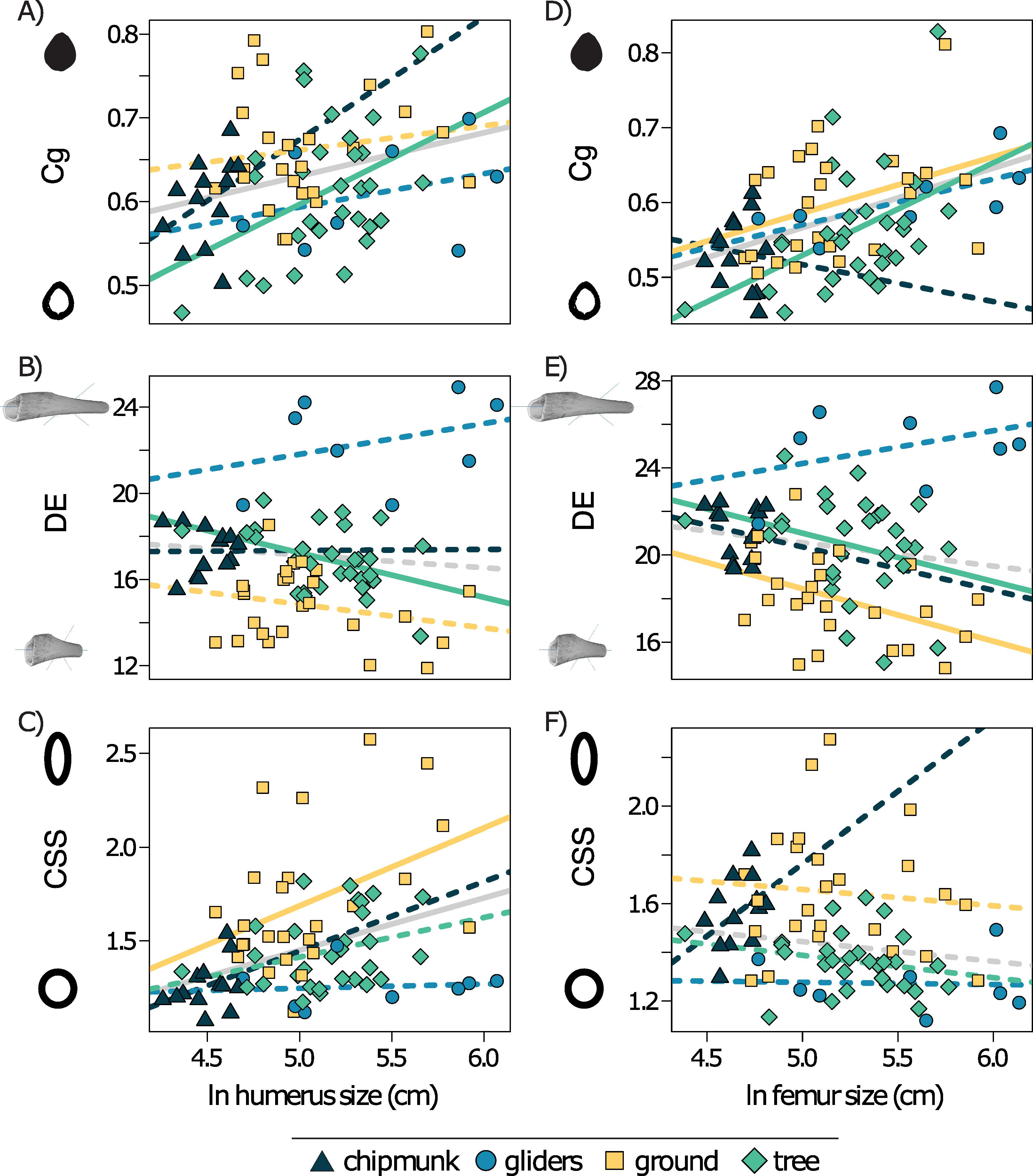
Scatter plot of ln limb size and each bone structure variable in the humerus (A-C) and femur (D-F). Confidence intervals that deviated from an isometric slope of 0 were interpreted as exhibiting significant positive allometry (slope > 0) or negative allometry (slope < 0). Solid lines depict significant allometry, whereas dashed lines depict no significant allometry. Gray lines indicate relationships across all squirrel ecotypes. Cg = global compactness; DE = diaphysis elongation; cross-sectional shape = CSS.

### Scaling and ecomorphology of limb bones within ecotypes

Scaling patterns in external shape and bone structure variables of the humerus and femur are further nuanced by ecological specialization. In ground squirrels, positive allometry towards more robust humeral shapes is consistent with positive allometry of more robust body shapes (Linden et al. 2023), which together may serve as adaptations for the digging and clearing stages of scratch digging during burrow construction. The digging stage consists of the claws of the forelimbs striking the ground while the hind limbs support the body (Gasc et al. 1985; Hildebrand 1995), and a plethora of skeletal and muscular adaptations in the forelimb are associated with this first stage (Lessa and Stein 1992; Hildebrand 1995; Lagaria and Youlatos 2006; Vassallo 2006; Steiner-Souza et al. 2010). Our finding of positive allometry in humeral shape and CSS provides additional evidence for adaptation towards digging, particularly in larger ground squirrels with more elliptical cross sections that would better resist uniaxial bending loads during digging. Trends towards negative allometry of humeral DE would also lead to more compact, robust humeri that could decrease bone strain during the digging stage. In contrast, humeral Cg scaled isometrically with size, indicating that compact humeri are important for digging across all ground squirrel sizes. These patterns are also consistent with the analyses of humeral shape and its phylomorphospace. Ground squirrel humeri occupied regions of phylomorphospace (+PC1) that are associated with more pronounced deltoid tuberosities (Fig. 4A) that would provide greater attachment surface area for deltoids and pectoral muscles, which are needed to increase the force output advantageous for digging (Hildebrand 1985; Samuels and Van Valkenburgh 2008). These humeri associated with +PC1 also exhibited more pronounced medial epicondyles that would provide greater attachment sites for the extensor group key for scratch digging (Samuels and Van Valkenburgh 2008; Steiner-Souza et al. 2010).

Unlike in the humerus, our results for the relationships between femoral morphology and digging behavior are not as consistent. The clearing stage of scratch digging consists of the hind limb removing excess soil through backward extension while the body is supported by the forelimbs (Gasc et al. 1985; Hildebrand 1995). Whereas some have hypothesized that more robust femora in fossorial species would improve stability and load transfer during clearing (Casinos et al. 1993; Hildebrand 1995), others have hypothesized that fossorial species exhibit less robust femora to reflect less rigidity in bending and torsion compared to species that jump or run with their hind limbs (Gambaryan 1974; Biknevicius 1993; Wölfer et al. 2019). Furthermore, some have also hypothesized that robust femora are not a necessary adaptation because the hind limbs are only used to shovel away excess substrate during the clearing stage that had previously been loosened with the forelimbs during the digging stage (Gambaryan 1974; Wölfer et al. 2019). Our findings support both these hypotheses: positive allometry of femoral Cg and negative allometry of DE indicates that ground squirrel femora evolved to be more compact and robust with increasing femoral size, which may decrease bone deformation during the clearing phase. Conversely, the lack of allometry in femoral shape and CSS suggests that the femur is less specialized for digging compared to the humerus. Ground squirrel femora occupied regions of phylomorphospace (+PC1) (Fig. 4B) that are associated with more pronounced greater trochanters. These adaptations reflect the larger muscle attachment area for the gluteal muscles, which when enlarged in fossorial species may help the body resist being pushed back when digging, respectively (Samuels and Van Valkenburgh 2008).

Tree squirrels exhibited positive allometry in humeral and femoral Cg with increasing bone size. Additionally, we found that tree squirrels evolved more robust diaphyses (i.e., lower DE) in both the humerus and femur with increasing bone size. This contrasts with prior findings that scansorial or arboreal mammals often exhibit more elongate limb bones as size increases to increase limb span (Polly 2007; Kilbourne and Hoffman 2013). However, this may be due to the somewhat unique climbing method employed by tree squirrels, which use their claws to interlock into tree bark and their muscle strength to hold their bodies close to the tree as they climb (Schmidt and Fischer 2011). Furthermore, allometric trends towards relatively shorter humeri may increase mechanical advantage during climbing, given that the elbow functions like a class II lever (Keith 1919; Fleagle 1977). At larger body sizes, tree squirrels may need to counter the additional body weight by remaining relatively closer to the tree to minimize opposing forces. No significant relationships were observed between either humeral or femoral CSS and bone size in tree squirrels. This may point to a more circular cross-sectional shapes across different sizes as important in resisting multidirectional bending while climbing and jumping between trees (Patel et al. 2013).

The external shape and structure of chipmunk humeri and femora were all isometric. The lack of allometry in chipmunks may be due to their small sizes and limited size range. Prior research in other small mammals have shown that the ability of bone tissue to remodel in response to its mechanical environment (i.e., Wolff’s Law) may be restricted by body size (Dawson 1980; Meier et al. 2013). This makes specialization of bone structure traits such as compactness less likely to occur, as overall humeral morphology may be sufficient for digging or climbing without further cortical adaptations. Another explanation is that chipmunks primarily dig shallow shelters instead of extensive burrow systems and thus may not require extreme shape specializations (Samuels and Van Valkenburgh 2008). That chipmunk humeri and femora share overlapping regions of phylomorphospace with both tree and ground squirrels (Fig. 4) support these hypotheses.

Gliding squirrels evolved more gracile humeral and femoral shapes with increasing size, but, interestingly, humeral and femoral DE were isometric. The lack of DE allometry in gliding squirrels may be due to wing loading (body mass / patagium area); larger gliding squirrels decrease the detrimental effects of higher wing loading by exhibiting a lower relative body weight instead of evolving relatively longer limbs compared to smaller gliding squirrels (Thorington and Heaney 1981). Nevertheless, gliding squirrels on average exhibited more elongate humeri and femora compared to other ecotypes (Fig. 4; Fig. 6), consistent with previous findings (Peterka 1936; Bryant 1945; Thorington and Heaney 1981; Grossnickle et al. 2020; Linden et al. 2023). Furthermore, gliding squirrels tended to exhibit the most circular cross sections of the humerus and femur (i.e., low CSS values; Fig. 6), which would maximize resistance of torsional stresses during aerial locomotion (Swartz et al. 1992) or like tree squirrels multidirectional bending while climbing and jumping between trees (Patel et al. 2013).

**Fig. 6.**
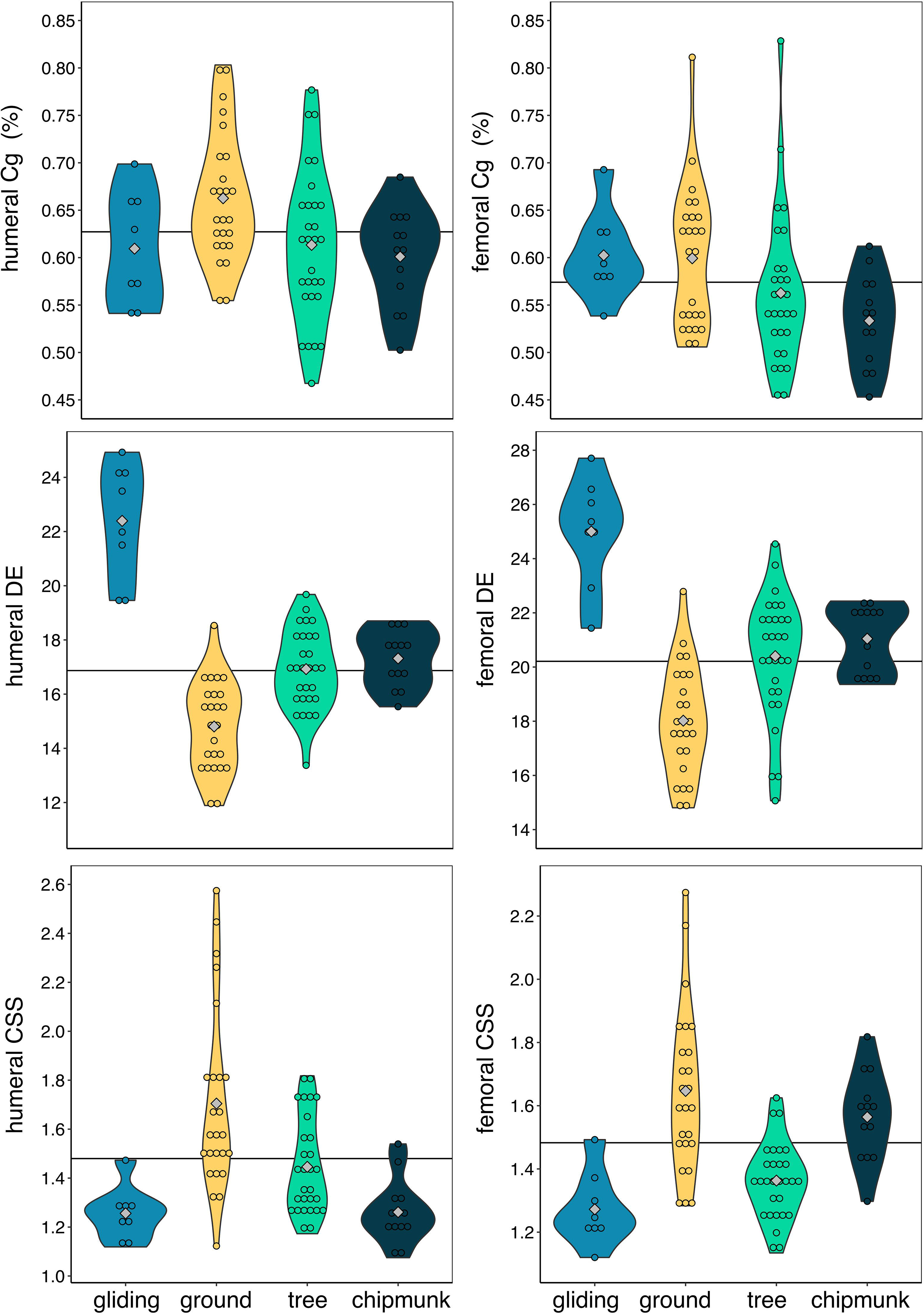
Violin plots of raw bone structure variables in humeri and femora. Variables are not corrected for allometry. The gray diamonds indicate the mean of each ecotype, and the horizontal black line indicates the mean size across all ecotypes. Cg = global compactness; DE = diaphysis elongation; cross-sectional shape = CSS.

### Comparing humeral and femoral shape and bone structure

Humeral shape and femoral shape are tightly correlated across squirrels (r-PLS = 0.91, Z = 5.884, P < 0.001), a pattern that is consistent with other mammals (e.g., carnivorans, Martín-Serra et al. 2015; Hanot et al. 2017; Hedrick et al. 2019). Together, this indicates that the relative robustness of the humerus and femur contributes the greatest covariation between the two bones. However, we found that these patterns differed between scansorial and fossorial ecotypes: while gliding squirrels and tree squirrels exhibit correlated humeral and femoral shapes, chipmunks and ground squirrels do not. These differences may not be surprising because the humerus and femur exhibit different functions during the digging and clearing stages, respectively, of burrow construction (see previous section).

Ecotype may also influence whether bone structure traits differ between the humerus and femur. Previous studies suggest that scansorial mammals exhibit similar global compactness values between the forelimb and the hindlimb due to the equal importance of both limbs during climbing, whereas scratch-digging mammals exhibit higher compactness in the forelimb due to the many specializations associated with digging (Hildebrand 1985; Straehl et al. 2013). We found higher mean global compactness in humeri than in femora in chipmunks and ground squirrels, an expected result since chipmunks and ground squirrels exhibit forelimb specializations associated with a fossorial lifestyle (Lessa and Stein 1992; Hildebrand 1995; Lagaria and Youlatos 2006; Vassallo 2006; Steiner-Souza et al. 2010). Surprisingly, tree squirrels also exhibited higher mean global compactness in humeri than in femora. A possible explanation is that although tree squirrels are scansorial, they are not limited to only climbing. Tree squirrels are often found running on the forest floor, and often bury food by scratch-digging (Ferron 1981). This may help to explain some of the adaptions seen in tree squirrels that have been historically correlated with fossorial locomotion. We also found higher DE values in the femur compared to the humerus in each of the four ecotypes. More elongate femora support previous findings that rodent humeri tend to be the shortest long bone whereas rodent femora are typically one of the longest (Prodinger et al. 2018). Lastly, CSS values do not differ between the humerus and femur, which is consistent with findings that humeral and femoral shape covary along the degree of bone robustness.

## Conclusion

We found that size, locomotor ecology, and underlying phylogenetic structure exhibit different relationships with the shape and structure of squirrel humeri and femora. Humeral and femoral shape is best predicted by ecotype, whereas humeral and femoral structure is best predicted by a combination of ecotype, size, and their interaction. Interestingly, the statistical relationships between these morphologies and ecotype were lost when accounting for phylogenetic relationships under Brownian motion. That all squirrel ecotypes are largely phylogenetically clustered invites questions about whether and how morphological variation in the humerus and femur evolved in relation to different ecologies early in squirrel evolutionary history and why they were subsequently maintained within clades. Clade-based evolutionary shifts in morphologies have been found in other mammals such as carnivorans (Law 2021; Law et al. 2022), and our results provide preliminary evidence that squirrels also exhibit clade-based evolutionary shifts in morphologies. Overall, these results indicate that mechanical and phylogenetic constraints and ecology may enact different pressures on the external and structural aspects of limb bone morphology. Increasing the number of species across the squirrel phylogeny and expanding to other rodents will enable us to tease apart the effects of selection and phylogeny on potentially converging limb morphologies. Together, this work provides a strong morphological foundation for future research investigating the evolutionary biomechanics and ecology of squirrel locomotion.

## Acknowledgements

The authors are grateful to the collections and its staff of the Burke Museum of Natural History, California Academy of Sciences, Florida Museum of Natural History, Museum of Vertebrate Zoology, Natural History Museum of Los Angeles County, Slater Museum of Natural History, Smithsonian Institute Natural Museum of Natural History, University of Kansas Biodiversity Institute & Natural History Museum, and Yale Peabody Museum. We thank Jonathan Huie for helpful suggestions on using SegmentGeometry, Kelly Diamond for helpful suggestions on using SlicerMorph, Jack Tseng and Dave Grossnickle for CT scanning some specimens, the staff of Friday Harbor Labs for allowing us to CT scan specimens, and Jan Wölfer and Jesse Young for their valuable reviews.

## Data Availability Statement

The data underlying this article are available in the Dryad Digital Repository, at https://dx.doi.org/[XXXXX]

## References

Adams D, Collyer M, Kaliontzopoulou A, Baken E. 2022. Geomorph: Software for geometric morphometric analyses.

Adams DC, Collyer ML. 2018. Multivariate Phylogenetic Comparative Methods: Evaluations, Comparisons, and Recommendations. Syst Biol 67:14–31.

Alexander RMcN, Jayes AS, Maloiy GMO, Wathuta EM. 1979. Allometry of the limb bones of mammals from shrews (Sorex) to elephant (Loxodonta). J Zool 189:305–14.

Amson E, Bibi F. 2021. Differing effects of size and lifestyle on bone structure in mammals. BMC Biol 19:87.

Amson E, Billet G, de Muizon C. 2018. Evolutionary adaptation to aquatic lifestyle in extinct sloths can lead to systemic alteration of bone structure. Proc R Soc B Biol Sci 285:20180270.

Amson E, Scheyer TM, Martinez Q, Schwermann AH, Koyabu D, He K, Ziegler R. 2022. Unique bone microanatomy reveals ancestry of subterranean specializations in mammals. Evol Lett 6:552–61.

Baken EK, Collyer ML, Kaliontzopoulou A, Adams DC. 2021. geomorph v4.0 and gmShiny: Enhanced analytics and a new graphical interface for a comprehensive morphometric experience. Methods Ecol Evol 12:2355–63.

Berman AG, Clauser CA, Wunderlin C, Hammond MA, Wallace JM. 2015. Structural and Mechanical Improvements to Bone Are Strain Dependent with Axial Compression of the Tibia in Female C57BL/6 Mice. Plos One 10:e0130504

Biewener AA. 1983. Allometry of Quadrupedal Locomotion: the Scaling of Duty Factor, Bonecurvature and Limb Orientation to Body Size. J Exp Biol 105:147–71.

Biewener AA. 2005. Biomechanical consequences of scaling. J Exp Biol 208:1665–1676.

Biewener AA. 1990. Biomechanics of Mammalian Terrestrial Locomotion. Science 250:1097–1103.

Biknevicius AR. 1993. Biomechanical Scaling of Limb Bones and Differential Limb Use in Caviomorph Rodents. J Mammal 74:95–107.

Bryant MD. 1945. Phylogeny of Nearctic Sciuridae. Am Midl Nat 33:257.

Burr DB, Ruff CB, Johnson C. 1989. Structural adaptations of the femur and humerus to arboreal and terrestrial environments in three species of macaque. Am J Phys Anthropol 79:357– 67.

Campione NE, Evans DC. 2012. A universal scaling relationship between body mass and proximal limb bone dimensions in quadrupedal terrestrial tetrapods. BMC Biol 10:60.

Casinos A, Quintana C, Viladiu C. 1993. Allometry and adaptation in the long bones of a digging group of rodents (Ctenomyinae). Zool J Linn Soc 107:107–15.

Christiansen P. 1999. Scaling of the limb long bones to body mass in terrestrial mammals. J Morphol 239:167–90.

Clavel J, Aristide L, Morlon H. 2019. A Penalized Likelihood Framework for High-Dimensional Phylogenetic Comparative Methods and an Application to New-World Monkeys Brain Evolution. Syst Biol 68:93–116.

Clavel J, Escarguel G, Merceron G. 2015. mvmorph: an r package for fitting multivariate evolutionary models to morphometric data. Methods Ecol Evol 6:1311–19.

Clavel J, Morlon H. 2020. Reliable Phylogenetic Regressions for Multivariate Comparative Data: Illustration with the MANOVA and Application to the Effect of Diet on Mandible Morphology in Phyllostomid Bats. Syst Biol 69:927–43.

Collyer ML and Adams DC. 2018. RRPP: An r package for fitting linear models to high_Jdimensional data using residual randomization. Methods in Ecology and Evolution, 9:1772–1779.

Collyer ML, Sekora DJ, Adams DC. 2015. A method for analysis of phenotypic change for phenotypes described by high-dimensional data. Heredity 115:357–65.

Cowin SC, Sadegh AM, Luo GM. 1992. An Evolutionary Wolff’s Law for Trabecular Architecture. J Biomech Eng 114:129–36.

Currey JD, Alexander RMcN. 1985. The thickness of the walls of tubular bones. J Zool 206:453–68.

Dawson DL. 1980. Functional interpretations of the radiographic anatomy of the femora of Myotis iucifugus, Pipistrellus subflavus, and Blarina brevicauda. Am J Anat 157:1–15.

Demes B, Jungers WL. 1993. Long bone cross-sectional dimensions, locomotor adaptations and body size in prosimian primates. J Hum Evol 25:57–74.

Diamond KM, Burtner AE, Siddiqui D, Alvarado K, Leake SL, Rolfe S, Zhang C, Kwon RY, Maga AM. 2022. Examining craniofacial variation among crispant and mutant zebrafish models of human skeletal diseases.

Doube M, Conroy AW, Christiansen P, Hutchinson JR, Shefelbine S. 2009. Three-Dimensional Geometric Analysis of Felid Limb Bone Allometry. PLOS ONE 4:e4742.

Etienne C, Filippo A, Cornette R, Houssaye A. 2021. Effect of mass and habitat on the shape of limb long bones: A morpho-functional investigation on Bovidae (Mammalia: Cetartiodactyla). J Anat 238:886–904.

Fabre A-C, Cornette R, Goswami A, Peigné S. 2015. Do constraints associated with the locomotor habitat drive the evolution of forelimb shape? A case study in musteloid carnivorans. J Anat 226:596–610.

Fabre A-C, Cornette R, Peigné S, Goswami A. 2013. Influence of body mass on the shape of forelimb in musteloid carnivorans. Biol J Linn Soc 110:91–103.

Fedorov A, Beichel R, Kalpathy-Cramer J, Finet J, Fillion-Robin J-C, Pujol S, Bauer C, Jennings D, Fennessy F, Sonka M, Buatti J, Aylward S, Miller JV, Pieper S, Kikinis R. 2012. 3D Slicer as an image computing platform for the Quantitative Imaging Network. Magn Reson Imaging, Quantitative Imaging in Cancer 30:1323–41.

Fleagle JG. 1977. Locomotor behavior and muscular anatomy of sympatric Malaysian leaf-monkeys (*Presbytis obscura* and *Presbytis melalophos*). Am J Phys Anthropol 46:297–307.

Ferron J. 1981. Comparative Ontogeny of Behaviour in Four Species of Squirrels (Sciuridae). Z Für Tierpsychol 55:193–216.

Gambaryan PP. 1974. How Mammals Run: Anatomical Adaptations Wiley.

Gasc J-P, Renous S, Casinos A, Laville E, Bou J. 1985. Comparison of diverse digging patterns in some small mammals. Fortschr Zool 30:35–38.

Grossnickle DM, Chen M, Wauer JGA, Pevsner SK, Weaver LN, Meng Q-J, Liu D, Zhang Y-G, Luo Z-X. 2020. Incomplete convergence of gliding mammal skeletons*. Evolution 74:2662–80.

Hanot P, Herrel A, Guintard C and Cornette R. 2017. Morphological integration in the appendicular skeleton of two domestic taxa: the horse and donkey. Proc. R. Soc. B.2842017124120171241

Hayssen V. 2008. Patterns of Body and Tail Length and Body Mass in Sciuridae. In: Journal of Mammalogy p. 852–73.

Hedrick BP, Dickson BV, Dumont ER, Pierce SE. 2020. The evolutionary diversity of locomotor innovation in rodents is not linked to proximal limb morphology. Scientific Reports, 1–11.

Hildebrand M. 1985. Chapter 6. Digging of Quadrupeds. In: Hildebrand M, Bramble DM, Liem KF, Wake DB, editors. Functional Vertebrate Morphology Harvard University Press. p. 89–109.

Hildebrand M. 1995. Analysis of vertebrate structure. 4th ed. ed New York: J. Wiley.

Huie JM, Summers AP, Kawano SM. 2022. SegmentGeometry: A Tool for Measuring Second Moment of Area in 3D Slicer. Integr Org Biol 4:obac009.

Hunt NH, Jinn J, Jacobs LF, Full RJ. 2021. Acrobatic squirrels learn to leap and land on tree branches without falling. Science 373:697–700.

Keith A. 1919. The Engines of the Human Body. London: Williams and Norgate.

Kikinis R, Pieper SD, Vosburgh KG. 2014. 3D Slicer: A Platform for Subject-Specific Image Analysis, Visualization, and Clinical Support. In: Jolesz FA, editor. Intraoperative Imaging and Image-Guided Therapy New York, NY: Springer. p. 277–89.

Kilbourne BM, Hutchinson JR. 2019. Morphological diversification of biomechanical traits: mustelid locomotor specializations and the macroevolution of long bone cross-sectional morphology. Bmc Evol Biol 19:37.

Kilbourne BM, Hoffman LC. 2013. Scale Effects between Body Size and Limb Design in Quadrupedal Mammals. PLOS ONE 8:e78392.

Kimura T. 1991. Long and robust bones of primates. In: Ehara A, editor. Primatology Today: Proceedings of the XIIIth Congress of the International Primatological Society, Nagoya and Kyoto, 18-24, July 1990 Amsterdam; New York: Elsevier Science Publishers.

Kivell TL. 2016. A review of trabecular bone functional adaptation: what have we learned from trabecular analyses in extant hominoids and what can we apply to fossils? J Anat 228:569–94.

Lagaria A, Youlatos D. 2006. Anatomical Correlates to Scratch Digging in the Forelimb of European Ground Squirrels (Spermophilus citellus). J Mammal 87:563–70.

Law CJ. 2021. Ecological Drivers of Carnivoran Body Shape Evolution. Am Nat 198:406–20.

Law CJ, Blackwell EA, Curtis AA, Dickinson E, Hartstone-Rose A, Santana SE. 2022. Decoupled evolution of the cranium and mandible in carnivoran mammals. Evolution 76:2959–74.

Lessa EP, Stein BR. 1992. Morphological constraints in the digging apparatus of pocket gophers (Mammalia: Geomyidae). Biol J Linn Soc 47:439–53.

Lieberman D, Polk J, and Demes B. 2004. Predicting long bone loading from cross-sectional geometry. Am J Phys Anthropol 123:156–171.

Linden T, Burtner A, Rickman J, McFeely A, Santana S, Law C. 2023. Scaling patterns of body plans differ among squirrel ecotypes. PeerJ 11:e14800.

Martín-Serra A, Figueirido B, Palmqvist P. 2014a. A Three-Dimensional Analysis of Morphological Evolution and Locomotor Performance of the Carnivoran Forelimb. PloS One 9:e85574.

Martín-Serra A, Figueirido B, Palmqvist P. 2014b. A three-dimensional analysis of the morphological evolution and locomotor behaviour of the carnivoran hind limb. BMC Evol Biol 14:129.

Martín_JSerra A, Figueirido B, Pérez_JClaros JA, and Palmqvist P. 2015. Patterns of morphological integration in the appendicular skeleton of mammalian carnivores. Evolution 69:321–340.

McMahon T. 1973. Size and shape in biology. Science 179:1201–4.

Meier PS, Bickelmann C, Scheyer TM, Koyabu D, Sánchez-Villagra MR. 2013. Evolution of bone compactness in extant and extinct moles (Talpidae): exploring humeral microstructure in small fossorial mammals. BMC Evol Biol 13:55.

Mielke M, Wölfer J, Arnold P, van Heteren AH, Amson E, Nyakatura JA. 2018. Trabecular architecture in the sciuromorph femoral head: allometry and functional adaptation. Zool Lett 4:10.

Montoya-Sanhueza G, Chinsamy A. 2017. Long bone histology of the subterranean rodent Bathyergus suillus (Bathyergidae): ontogenetic pattern of cortical bone thickening. J Anat 230:203–33.

Patel BA, Ruff CB, Simons ELR, Organ JM. 2013. Humeral cross-sectional shape in suspensory primates and sloths. Anat Rec. 556:545–56.

Peterka HE. 1936. A Study of the Myology and Osteology of Tree Sciurids with Regard to Adaptation to Arboreal, Glissant and Fossorial Habits. Trans Kans Acad Sci 1903–39:313–32.

Polly PD. 2007. Limbs in Mammalian Evolution. In: Hall KB, editor. Fins into Limbs: Evolution, Development, and Transformation Chicago: University of Chicago Press. p. 245–68.

Porto A, Rolfe S, Maga AM. 2021. ALPACA: A fast and accurate computer vision approach for automated landmarking of three-dimensional biological structures. Methods Ecol Evol 12:2129–44.

Prodinger PM, Foehr P, Bürklein D, Bissinger O, Pilge H, Kreutzer K, von Eisenhart-Rothe R, Tischer T. 2018. Whole bone testing in small animals: systematic characterization of the mechanical properties of different rodent bones available for rat fracture models. Eur J Med Res 23:8.

Rohlf FJ, Slice D. 1990. Extensions of the Procrustes Method for the Optimal Superimposition of Landmarks. Syst Biol 39:40–59.

Rolfe S, Pieper S, Porto A, Diamond K, Winchester J, Shan S, Kirveslahti H, Boyer D, Summers A, Maga AM. 2021. SlicerMorph: An open and extensible platform to retrieve, visualize and analyse 3D morphology. Methods Ecol Evol 12:1816–25.

Ruff C, Holt B, Trinkaus E. 2006. Who’s afraid of the big bad Wolff?: “Wolff’s law” and bone functional adaptation. Am J Phys Anthropol 129:484–98.

Ryan TM, Shaw CN. 2013. Trabecular bone microstructure scales allometrically in the primate humerus and femur. Proc R Soc B Biol Sci 280:20130172.

Samuels JX, Van Valkenburgh B. 2008. Skeletal indicators of locomotor adaptations in living and extinct rodents. J Morphol 269:1387–1411.

Schaffler MB, Burr DB, Jungers WL, Ruff CB. 1985. Structural and Mechanical Indicators of Limb Specialization in Primates. Folia Primatol (Basel) 45:61–75.

Scheidt A, Wölfer J, Nyakatura JA. 2019. The evolution of femoral cross-sectional properties in sciuromorph rodents: Influence of body mass and locomotor ecology. J Morphol 280:1156–69.

Schlager S. 2017. Chapter 9 - Morpho and Rvcg – Shape Analysis in R: R-Packages for Geometric Morphometrics, Shape Analysis and Surface Manipulations. In: Zheng G, Li S, Székely G, editors. Statistical Shape and Deformation Analysis Academic Press. p. 217–56.

Schmidt A, Fischer MS. 2011. The kinematic consequences of locomotion on sloped arboreal substrates in a generalized (Rattus norvegicus) and a specialized (Sciurus vulgaris) rodent. J Exp Biol 214:2544–59.

Smith JM, Savage RJG. 1956. Some locomotory adaptations in mammals. J Linn Soc Lond Zoology 42:603–22.

Steiner-Souza F, De Freitas TRO, Cordeiro-Estrela P. 2010. Inferring adaptation within shape diversity of the humerus of subterranean rodent Ctenomys. Biol J Linn Soc 100:353–67.

Straehl FR, Scheyer TM, Forasiepi AM, MacPhee RD, Sánchez-Villagra MR. 2013. Evolutionary Patterns of Bone Histology and Bone Compactness in Xenarthran Mammal Long Bones. PLOS ONE 8:e69275.

Swartz SM, Bennett MB, Carrier DR. 1992. Wing bone stresses in free flying bats and the evolution of skeletal design for flight. Nature 359:726–29.

Thorington RW Jr, Heaney LR. 1981. Body Proportions and Gliding Adaptations of Flying Squirrels (Petauristinae). J Mammal 62:101–14.

Tung Ho L si, Ané C. 2014. A Linear-Time Algorithm for Gaussian and Non-Gaussian Trait Evolution Models. Syst Biol 63:397–408.

Upham NS, Esselstyn JA, Jetz W. 2019. Inferring the mammal tree: Species-level sets of phylogenies for questions in ecology, evolution, and conservation. PLOS Biol 17:e3000494.

Vassallo A. 2006. Functional morphology, comparative behaviour, and adaptation in two sympatric subterranean rodents genus Ctenomys (Caviomorpha: Octodontidae). J Zool 244:415–27.

Wölfer J, Amson E, Arnold P, Botton-Divet L, Fabre A-C, van Heteren AH, Nyakatura JA. 2019. Femoral morphology of sciuromorph rodents in light of scaling and locomotor ecology. J Anat 234:731–47.

Zelditch ML, Swiderski DL, Sheets HD. 2012. Chapter 1 - Introduction. In: Zelditch ML, Swiderski DL, Sheets HD, editors. Geometric Morphometrics for Biologists (Second Edition) San Diego: Academic Press. p. 1–20.

