## Supplementary Materials for "Size and locomotor ecology have differing effects on the external and internal morphologies of squirrel (Rodentia: Sciuridae) limb bones"

Table S1. Museum specimens used in this study. CAS = California Academy of Sciences; KU = University of Kansas Biodiversity Institute & Natural History Museum; LACM = Natural History Museum of Los Angeles County; MVZ = UC Berkeley Museum of Vertebrate Zoology; PSM = University of Puget Sound Slater Museum of Natural History; UF = Florida Museum of Natural History; USNM = Smithsonian National Museum of Natural History; UWBM = University of Washington Burke Museum of Natural History

| species | catalog number | ecotype |
| --- | --- | --- |
| *Ammospermophilus leucurus* | UWBM74646 | ground |
| *Ammospermophilus nelsoni* | MVZ234368 | ground |
| *Callosciurus erythraeus* | CAS6371 | tree |
| *Callosciurus prevostii* | LACM97346 | tree |
| *Callosciurus pygerythrus* | PSM27419 | tree |
| *Callospermophilus lateralis* | UWBM38960 | ground |
| *Callospermophilus madrensis* | MVZ99799 | ground |
| *Callospermophilus saturatus* | UWBM38984 | ground |
| *Cynomys gunnisoni* | MVZ99763 | ground |
| *Cynomys ludovicianus* | UWBM75774 | ground |
| *Eoglaucomys fimbriatus* | USNM173363 | gliding |
| *Funambulus palmarum* | MVZ183704 | tree |
| *Funambulus pennantii* | UF30290 | tree |
| *Funisciurus pyrropus* | MVZ196233 | tree |
| *Glaucomys sabrinus* | UWBM35058 | gliding |
| *Glaucomys volans* | UWBM43897 | gliding |
| *Heliosciurus mutabilis* | MVZ220919 | tree |
| *Heliosciurus rufobrachium* | MVZ196227 | tree |
| *Ictidomys mexicanus* | MVZ93789 | ground |
| *Iomys horsfieldii* | PSM10370 | gliding |
| *Marmota broweri* | UWBM82831 | ground |
| *Marmota caligata* | UWBM31095 | ground |
| *Marmota flaviventris* | UWBM32500 | ground |
| *Marmota monax* | UWBM39792 | ground |
| *Microsciurus flaviventer* | MVZ154935 | tree |
| *Otospermophilus variegatus* | UWBM79876 | ground |
| *Petaurista alborufus* | MVZ183717 | gliding |
| *Petaurista petaurista* | USNM197320 | gliding |
| *Petaurista philippensis* | USNM314973 | gliding |
| *Poliocitellus franklinii* | UWBM33260 | ground |
| *Protoxerus stangeri* | MVZ196230 | tree |
| *Pteromys volans* | UWBM39689 | gliding |
| *Ratufa bicolor* | MVZ123699 | tree |
| *Ratufa macroura* | UWBM21183 | tree |
| *Sciurus aberti* | UWBM35277 | tree |
| *Sciurus aestuans* | MVZ182070 | tree |
| *Sciurus aureogaster* | LACM053653 | tree |
| *Sciurus colliaei* | LACM058790 | tree |
| *Sciurus deppei* | MVZ98319 | tree |
| *Sciurus griseus* | UWBM76257 | tree |
| *Sciurus nayaritensis* | UWBM81981 | tree |
| *Sciurus niger* | UWBM35230 | tree |
| *Sciurus spadiceus* | MVZ166023 | tree |
| *Sciurus stramineus* | MVZ135638 | tree |
| *Sciurus variegatoides* | MVZ131091 | tree |
| *Sciurus vulgaris* | UWBM39069 | tree |
| *Sundasciurus juvencus* | KU165382 | tree |
| *Tamias alpinus* | MVZ207209 | chipmunk |
| *Tamias amoenus* | UWBM31698 | chipmunk |
| *Tamias cinereicollis* | UWBM38257 | chipmunk |
| *Tamias dorsalis* | UWBM78908 | chipmunk |
| *Tamias merriami* | UWBM60172 | chipmunk |
| *Tamias minimus* | UWBM30813 | chipmunk |
| *Tamias quadrivittatus* | UWBM35301 | chipmunk |
| *Tamias ruficaudus* | UWBM34354 | chipmunk |
| *Tamias senex* | UWBM78783 | chipmunk |
| *Tamias sibiricus* | UWBM39259 | chipmunk |
| *Tamias speciosus* | UWBM43059 | chipmunk |
| *Tamias striatus* | UWBM35246 | chipmunk |
| *Tamias townsendii* | UWBM35213 | chipmunk |
| *Tamiasciurus douglasii* | UWBM81947 | tree |
| *Tamiasciurus hudsonicus* | UWBM81510 | tree |
| *Tamiops maritimus* | MVZ186481 | tree |
| *Tamiops swinhoei* | UWBM75283 | tree |
| *Urocitellus armatus* | MVZ64641 | ground |
| *Urocitellus beldingi* | UWBM42459 | ground |
| *Urocitellus columbianus* | UWBM74770 | ground |
| *Urocitellus elegans* | UWBM33286 | ground |
| *Urocitellus parryii* | UWBM39268 | ground |
| *Urocitellus richardsonii* | UWBM32960 | ground |
| *Urocitellus townsendii* | UWBM78333 | ground |
| *Xerospermophilus spilosoma* | UWBM35284 | ground |
| *Xerospermophilus tereticaudus* | UWBM35502 | ground |
| *Xerus erythropus* | MVZ115437 | ground |
| *Xerus inauris* | MVZ117287 | ground |

Table S2. The observed difference between the mean coefficients of ecotype pairs and its 95% confidence interval. Confidence intervals that encompassed zero indicated that ecotype pairs were not significantly different from each other.

| Humerus | |  |  |  |
| --- | --- | --- | --- | --- |
| differences between mean humeral size | | |  |  |
|  |  | chipmunk | gliding | ground |
|  | gliding | 0.82 [0.79:0.85] |  |  |
|  | ground | 0.40 [0.36:0.43] | 0.42 [0.40:0.45] |  |
|  | tree | 0.61 [0.59:0.64] | 0.21 [0.18:0.23] | 0.22 [0.19:0.24] |
| difference between mean slopes for humeral Cg scaling patterns | | | | |
|  |  | chipmunk | gliding | ground |
|  | gliding | 0.11 [0.10:0.12] |  |  |
|  | ground | 0.12 [0.11:0.13] | 0.01 [0.01:0.02] |  |
|  | tree | 0.04 [0.03:0.05] | 0.07 [0.06:0.08] | 0.08 [0.08:0.09] |
| differences between mean humeral Cg | | |  |  |
|  |  | chipmunk | gliding | ground |
|  | gliding | 0.02 [0.01:0.04] |  |  |
|  | ground | 0.07 [0.06:0.08] | 0.04 [0.03:0.05] |  |
|  | tree | 0.03 [0.02:0.04] | 0.01 [0.00:0.02] | 0.03 [0.03:0.04] |
| difference between mean slopes for humerus DE scaling patterns | | | | |
|  |  | chipmunk | gliding | ground |
|  | gliding | 1.26 [1.05:1.46] |  |  |
|  | ground | 1.28 [1.08:1.48] | 2.54 [2.45:2.62] |  |
|  | tree | 2.28 [2.08:2.48] | 3.53 [3.44:3.62] | 1.00 [0.92:1.07] |
| differences between mean humeral DE | | |  |  |
|  |  | chipmunk | gliding | ground |
|  | gliding | 5.06 [5.02:5.10] |  |  |
|  | ground | 2.52 [2.48:2.55] | 7.58 [7.54:7.62] |  |
|  | tree | 0.40 [0.37:0.43] | 5.46 [5.42:5.50] | 2.12 [2.09:2.14] |
| difference between mean slopes for humerus CSS scaling patterns | | | | |
|  |  | chipmunk | gliding | ground |
|  | gliding | 0.35 [0.32:0.39] |  |  |
|  | ground | 0.04 [0.01:0.07] | 0.39 [0.38:0.40] |  |
|  | tree | 0.16 [0.12:0.19] | 0.19 [0.18:0.21] | 0.20 [0.19:0.21] |
| differences between mean humeral CSS | | | |  |
|  |  | chipmunk | gliding | ground |
|  | gliding | 0.00 [0.00:0.01] |  |  |
|  | ground | 0.44 [0.44:0.45] | 0.45 [0.44:0.45] |  |
|  | tree | 0.19 [0.18:0.19] | 0.19 [0.18:0.19] | 0.26 [0.25:0.26] |
| Femur |  |  |  |  |
| differences between mean femoral size | | |  |  |
|  |  | chipmunk | gliding | ground |
|  | gliding | 0.79 [0.76:0.83] |  |  |
|  | ground | 0.48 [0.45:0.52] | 0.31 [0.28:0.34] |  |
|  | tree | 0.58 [0.55:0.61] | 0.21 [0.18:0.24] | 0.10 [0.07:0.12] |
| difference between mean slopes for femoral Cg scaling patterns | | | | |
|  |  | chipmunk | gliding | ground |
|  | gliding | 0.11 [0.10:0.12] |  |  |
|  | ground | 0.12 [0.11:0.14] | 0.01 [0.01:0.02] |  |
|  | tree | 0.17 [0.16:0.18] | 0.06 [0.06:0.07] | 0.05 [0.04:0.05] |
| differences between mean femoral Cg | | |  |  |
|  |  | chipmunk | gliding | ground |
|  | gliding | 0.08 [0.06:0.09] |  |  |
|  | ground | 0.08 [0.07:0.09] | 0.00 [-0.01:0.01] |  |
|  | tree | 0.05 [0.04:0.06] | 0.03 [0.02:0.04] | 0.03 [0.02:0.04] |
| difference between mean slopes for femoral DE scaling patterns | | | | |
|  |  | chipmunk | gliding | ground |
|  | gliding | 3.53 [3.18:3.88] |  |  |
|  | ground | 0.34 [0.00:0.68] | 3.87 [3.76:3.97] |  |
|  | tree | 0.15 [-0.19:0.50] | 3.68 [3.57:3.79] | 0.19 [0.09:0.28] |
| differences between mean femoral DE | | |  |  |
|  |  | chipmunk | gliding | ground |
|  | gliding | 3.95 [3.90:4.01] |  |  |
|  | ground | 3.05 [3.01:3.09] | 7.00 [6.95:7.05] |  |
|  | tree | 0.65 [0.61:0.69] | 4.60 [4.55:4.65] | 2.40 [2.36:2.43] |
| difference between mean slopes for femoral CSS scaling patterns | | | | |
|  |  | chipmunk | gliding | ground |
|  | gliding | 0.61 [0.58:0.64] |  |  |
|  | ground | 0.67 [0.64:0.70] | 0.07 [0.06:0.08] |  |
|  | tree | 0.70 [0.67:0.73] | 0.09 [0.08:0.10] | 0.02 [0.02:0.03] |
| differences between mean femoral CSS | | |  |  |
|  |  | chipmunk | gliding | ground |
|  | gliding | 0.29 [0.29:0.30] |  |  |
|  | ground | 0.08 [0.08:0.09] | 0.37 [0.37:0.38] |  |
|  | tree | 0.20 [0.20:0.20] | 0.09 [0.09:0.10] | 0.28 [0.28:0.29] |

Table S3. Results of two-block partial least squares (PLS) analyses showing relationships between limb bone shape and each bone structure trait of the humerus and femur.

|  |  | r-PLS | Z | P |  |  |  | r-PLS | Z | P |
| --- | --- | --- | --- | --- | --- | --- | --- | --- | --- | --- |
| humeral Cg vs shape | | |  |  |  | femoral Cg vs shape | | |  |  |
|  | all | 0.39 | 0.96 | 0.186 |  |  | all | 0.58 | 2.81 | **0.001** |
|  | chipmunk | 0.80 | 1.29 | 0.108 |  |  | chipmunk | 0.69 | -0.33 | 0.626 |
|  | gliding | 0.79 | -0.25 | 0.607 |  |  | gliding | 0.67 | -0.46 | 0.665 |
|  | ground | 0.73 | 1.38 | 0.089 |  |  | ground | 0.79 | 2.26 | **0.009** |
|  | tree | 0.74 | 1.94 | **0.028** |  |  | tree | 0.61 | 0.35 | 0.360 |
|  | humeral DE vs shape | | |  |  | femoral DE vs shape | | | |  |
|  | all | 0.88 | 4.55 | **0.001** |  |  | all | 0.85 | 4.70 | **0.001** |
|  | chipmunk | 0.80 | 1.34 | 0.100 |  |  | chipmunk | 0.71 | 0.00 | 0.493 |
|  | gliding | 0.82 | 0.17 | 0.438 |  |  | gliding | 0.72 | 0.05 | 0.467 |
|  | ground | 0.87 | 3.45 | **0.001** |  |  | ground | 0.78 | 2.24 | **0.011** |
|  | tree | 0.74 | 1.90 | **0.032** |  |  | tree | 0.85 | 3.30 | **0.001** |
|  | humeral CSS vs shape | | |  |  | femoral CSS vs shape | | | |  |
|  | all | 0.62 | 3.26 | **0.001** |  |  | all | 0.63 | 3.13 | **0.001** |
|  | chipmunk | 0.67 | -0.10 | 0.533 |  |  | chipmunk | 0.66 | -0.76 | 0.764 |
|  | gliding | 0.86 | 0.73 | 0.246 |  |  | gliding | 0.89 | 1.56 | 0.061 |
|  | ground | 0.76 | 1.91 | **0.025** |  |  | ground | 0.63 | 0.04 | 0.484 |
|  | tree | 0.76 | 2.15 | **0.015** |  |  | tree | 0.63 | 0.51 | 0.304 |

Table S4. Results of phylogenetic t-tests showing if each bone structure trait differed between the humerus and femur.

|  | mean  humeral Cg | mean  femoral Cg | phylogenetic  mean difference | Pagel’s λ | P |
| --- | --- | --- | --- | --- | --- |
| all squirrels | 0.63 | 0.57 | 0.05 | 0.10 | **0.001** |
| chipmunk | 0.60 | 0.53 | 0.13 | 0.91 | **0.043** |
| gliding | 0.61 | 0.60 | 0.01 | 0.00 | 0.595 |
| ground | 0.66 | 0.60 | 0.06 | 0.00 | **0.001** |
| tree | 0.61 | 0.56 | 0.05 | 0.52 | **0.010** |
|  | mean  humeral DE | mean  femoral DE | phylogenetic  mean difference | Pagel’s λ | P |
| all squirrels | 16.87 | 20.21 | -3.34 | 0.00 | **0.001** |
| chipmunk | 17.32 | 21.05 | -3.73 | 0.00 | **0.001** |
| gliding | 22.39 | 25.00 | -2.61 | 0.00 | **0.001** |
| ground | 14.81 | 18.02 | -3.21 | 0.00 | **0.001** |
| tree | 16.92 | 20.40 | -3.48 | 0.00 | **0.001** |
|  | mean  humeral CSS | mean  femoral CSS | phylogenetic  mean difference | Pagel’s λ | P |
| all squirrels | 1.48 | 1.48 | 0.06 | 0.36 | 0.531 |
| chipmunk | 1.26 | 1.56 | -0.30 | 0.00 | **0.001** |
| gliding | 1.26 | 1.27 | -0.02 | 0.00 | 0.733 |
| ground | 1.71 | 1.65 | 0.06 | 0.00 | 0.482 |
| tree | 1.45 | 1.36 | 0.08 | 0.00 | 0.065 |
